# Adolescent social profiles predict adult social behaviour, monoamine brain content and regional gene expression

**DOI:** 10.1101/2025.10.23.684125

**Authors:** N. Mirofle, S. Jamain, F. Chauveau, M. Penot, D. Chater, V. Latapie, T. Deneux, A. Faure, S. Granon

## Abstract

The study of inter-individual variability in animal behaviors during cognitive processes is crucial to better grasp the emergence of personality traits in healthy animals. Social experiences occurring at adolescence can affect the expression of the behavioural variability as it is a critical period of brain and cognitive plasticity. Here, we asked how individual variability of social profiles is shaped at adolescence and whether the profiles are associated with regional brain metabolites and gene expressions in the adult brain. At adolescence we determined a ‘dominant’ profile expressing more dominance, a ‘submissive’ profile expressing more submission and an ‘explorer’ profile showing more novelty exploration than social interaction. Adolescent social behaviors are conserved at adulthood. Our data suggest that dominance tendencies already present at adolescence are likely to stabilize throughout life, thus shaping future social behaviors. Our results show that social profiles are not related to sex and life environment. However, being more dominant, interactive and socially motivated shape dopaminergic olfactory activity at adolescence. Also, showing the explorer or submissive profiles is associated with an opposite serotonin pattern in the cerebellum, but a similar kynurenine pattern in the prefrontal cortex and cerebellum, thus pointing toward major neurotransmitter systems in key regions for brain health. This work opens new perspectives on the importance of studying social profiles early in life and the individual neurobiological associated features that could potentially predict individual vulnerability.

**Graphical Abstract:** 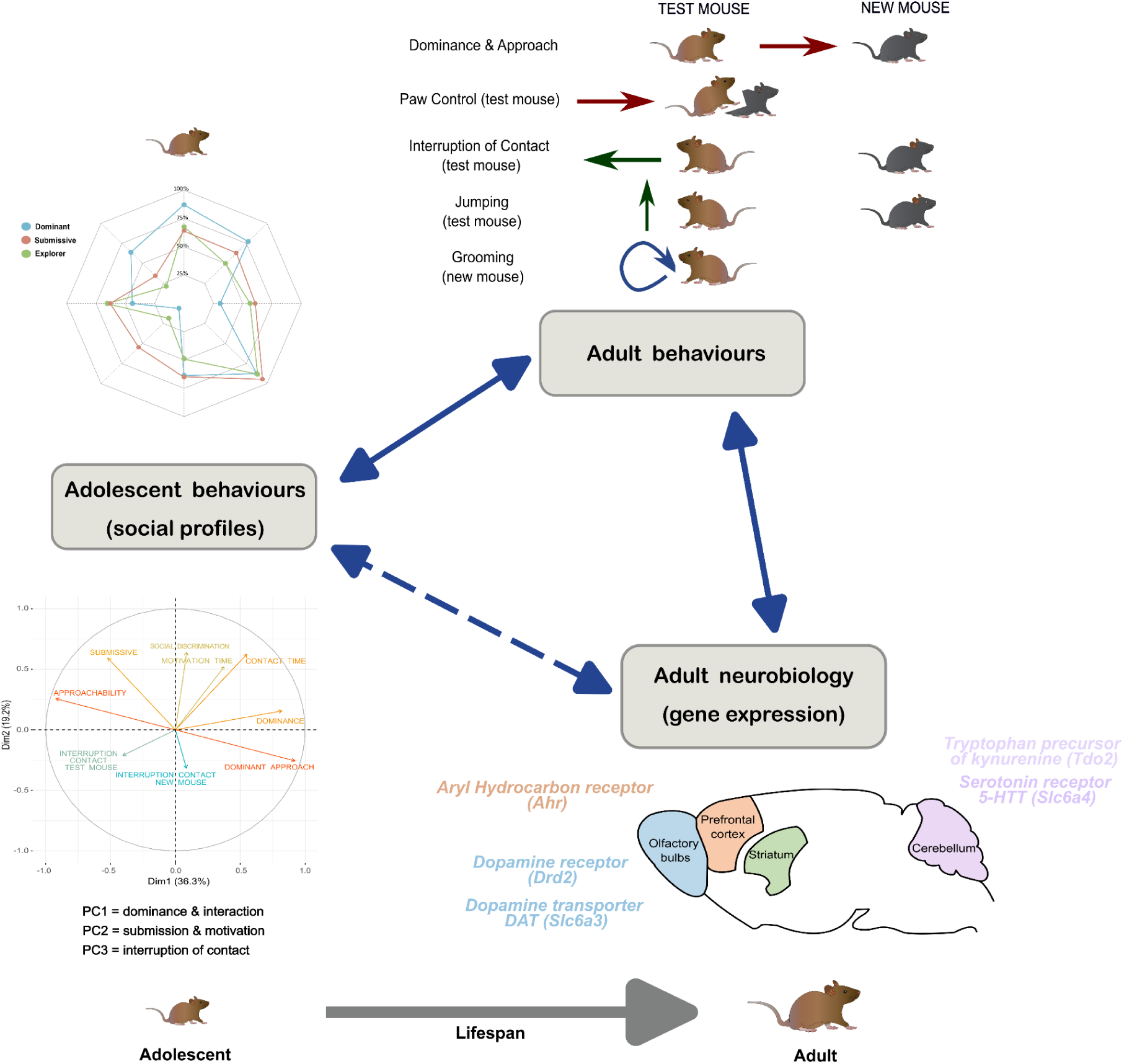

## Introduction

Individual variation in animal behaviors emerges in problem solving or development of cognitive strategies. This suggests the existence of animal personality, temperament, coping styles, or behavioural syndromes, terms that are not strictly equivalent but often used to describe similar ideas (reviews Dall, Houston & McNamara, 2004; Koolhaas et al., 2007; Réale et al., 2007). Inter-individual differences in physiology and behaviour observed in animal or human populations can reflect -and promote-individual’s adaptive capacity to provide appropriate responses to challenging or novel situations. Animal personality is a term borrowed from psychology. It refers to behavioural tendencies that differ across individuals, but remain consistent within the same individual over time and in different contexts (Réale et al., 2007). To that regard, studying individual variability in a specific context may offer a window into personality traits.

Salient experiences that occur at adolescence, *i.e*., after weaning, can affect the differential consistency of personality traits (review in Stamps & Groothuis 2010). Adolescence corresponds to a critical period of brain plasticity where the brain is more sensitive and influenced by experience (Peters & Naneix, 2022; Mancini et al., 2023), which seems to represent a favourable period for the emergence of behavioural variability.

Most mammals, including humans, are social species that develop, mature and evolve in a social environment. This development in the social environment may influence and shape both adaptive behaviors (Baarendse et al., 2013; Meyza et al., 2017) and their underlying neurobiological structures, particularly through epigenetic mechanisms (Becker et al., 2023). Social environments are therefore likely to have the strongest influence on behavior and brain plasticity (Baarendse et al., 2013). Indeed, early social experience is associated with neuronal and behavioural maturation (Baarendse et al., 2013; Mancini et al., 2023) in brain network and neurotransmitter systems common to social and cognitive behaviors. Such networks includes the prefrontal cortex, the cerebellum, the olfactory bulb, and striatal regions (Frith & Singer, 2008; Gangopadhyay et al., 2021; Duerler et al., 2022; Lanooij et al., 2023). Regarding neurotransmission systems, both monoamines (*i.e.* dopamine, serotonin and noradrenaline)(Pittaras et al., 2016; Fitoussi et al., 2018; Lanooij et al., 2023) and cholinergic projections (Lhopitallier et al., 2022; Caruso et al., 2018; Coura et al., 2013; Avale et al., 2011), along with the excitatory vs. inhibitory balance (*i.e.* GABA and Glutamate) (Caballero et al., 2016) were found to be crucial to adapt in social environments. Depending on the valence of early social experience (*i.e.* positive or negative), it can have long-term consequences through modulation of the kynurenin pathway (Dimonte et al., 2023), the oxytocin and vasopressin systems (Martucci et al., 2023; Shamay-Tsoory et al., 2009) and the expression of plasticity factors or cell cycle activity (Lopes et al., 2023). Previous studies have identified social decision profiles with specific social features associated with gene expression markers, and that remain stable throughout life (Forkosh et al., 2019; 2021).

Here, we asked how individual variability is shaped at adolescence, whether we can draw social profiles at that stage and at adulthood, and whether social profiles may be stable in life. We then asked if adolescent social profiles are associated with specific regional neurobiological -in the previous structures mentioned- and gene expression markers by focusing for the previous systems mentioned at the level of receptors and transporters. Finally we wondered whether we could predict different developmental trajectories for distinct early social profiles. In order to maximize individual development and to mimic heterogeneous life of healthy populations, we varied life environments (enriched vs. standard) and sex (male vs. female) and measured social and non social behaviors in a long term longitudinal study spanning from adolescence to adulthood.

## Results

### Interindividual differences in social behaviours at adolescence

To reveal social profiles at adolescence we focused on social choice when facing a new conspecific using two complementary social tasks: the social interaction task (SIT) and a social motivation task (3-chamber test, ‘3-ch’), which allowed us to collect numerous social parameters (interaction, contact escape, dominance, aggressiveness, communication), but also non-social parameters (novelty exploration, anxiety-like behaviour). During these tasks, the animal was free to interact or not with the new congener (of same sex and age) in a novel environment. Interindividual variability concerning all social parameters in both social tasks was studied using principal component analysis (PCA, **Figure 1** and **Supplementary Figure 1**), thus reducing variables into dimensions that are combining the most orthogonal ones best explaining the sample’s variance. PCA allowed comparison of individuals and identified the most relevant variables for classifying them. To prevent bias, only behaviours expressed by all animals were included (see **Figures 1** and **2**). Behaviours not expressed by all individuals (e.g. aggressiveness, ultrasonic communication) were not included in this analysis but analyzed elsewhere (**Supplementary Table 1**).

**Figure 1.**
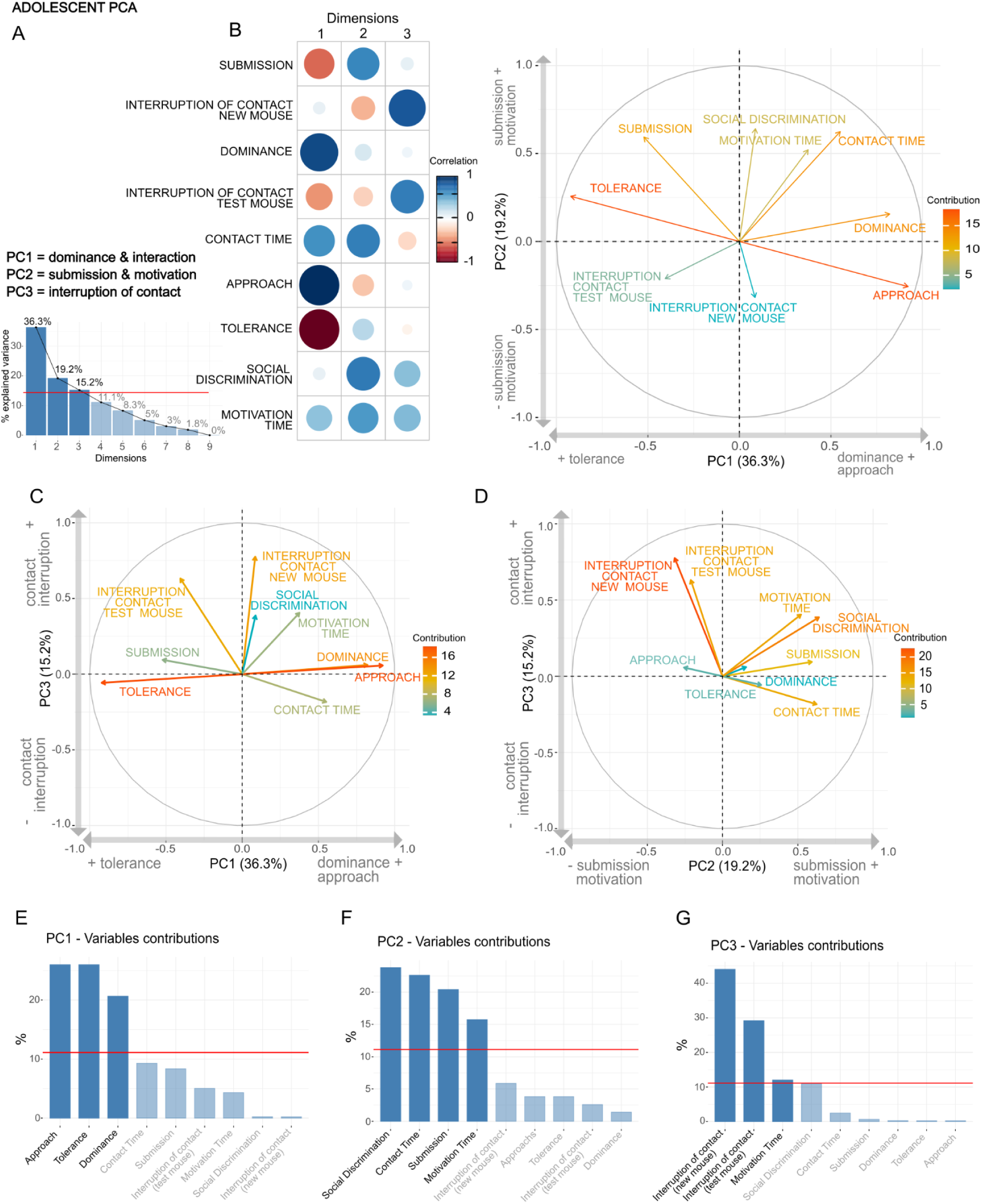
Identification of social profiles at adolescence. **A.** Description of the three PCA dimensions and each variable’s contribution to the total variance explained. **B.** Correlation between variables for all dimensions (left) and representation of the two first dimensions of the PCA analysis (right) **C.** PCA analysis for dimensions 1 and 3, and **D.** for dimensions 2 and 3. **E-G.** Detailed contribution of each variable in **E.** dimension 1, **F.** dimension 2 and **G.** dimension 3 separately. All PCA outputs are detailed in **Supplementary Table 1**. All variables were significantly contributing to the PCA analysis.

**Figure 2.**
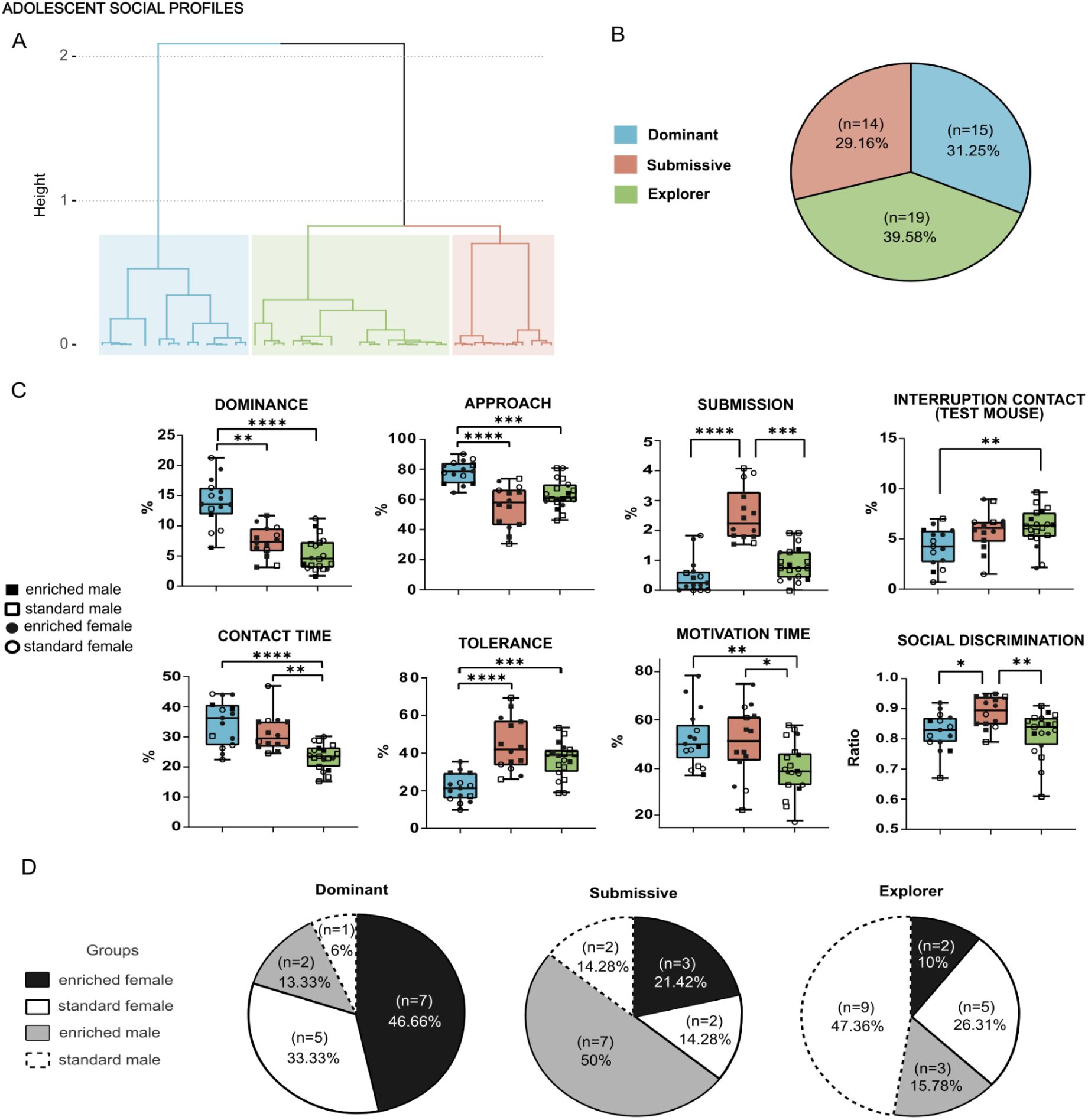
Characterisation of social profiles at adolescence. **A.** Hierarchical Clustering on Principal Components Analysis. **B.** Detailed individual distribution of animals in each profile. **C.** Statistical differences for behaviours characterising adolescent social profiles. **D.** Distribution by sex and life-environment for each profile. All variables were significantly different between profiles and were analysed with Kruskall-Wallis tests. * p < 0.05; ** p < 0.01; *** p < 0.001; **** p < 0.0001. Non-significant data, all pair-by-pair Dunn’s comparisons and other behaviours that were not included in this analysis (*i.e.*, aggressiveness, communication, exploration) are detailed in **Supplementary Table 1**.

PCA results revealed that 70.7% of the sample’s variance of social behaviour displayed at adolescence can be explained by 3 dimensions, as illustrated on **Figure 1 A.** Dimension 1 (PC1) was mainly composed of *interactions between mice* (approach from the host or from the new mice) and *dominance* (pursuit of the host mouse). Dimension 2 (PC2) was mostly constituted of *social motivation* (social discrimination in the 3-ch, contact time during SIT) and *submission* (host mouse followed by new mouse). Dimension 3 (PC3) consisted mainly in *interruption of contacts* and *motivation* (motivation time, *i.e.* contact time with social jar during 3-ch). We performed a Hierarchical Clustering on Principal Components (HCPC) analysis to identify specific groups according to their social dimensions (**Figure 2** and **Supplementary Figure 2**). We thus defined three groups to better capture interindividual variability regarding social behaviours at adolescence (**Supplementary Figure 2**). The three groups differed significantly for PC1 (Levene’s test: PC1: F(2, 45) = 3.7966, p = 0.03), PC2 (F(2,45) = 3.3313, p = 0.04), but not PC3 (F(2, 45) = 0.6878, p = 0.51).

To check whether this variability in social behaviour could be attributed to sex and environment alone, we plotted the different groups in the PCA analysis. Results illustrated on **Supplementary Figure 1** show that sex and environment independent factors did not solely explain the variance of the data while social profiles did. Indeed, while PCA outputs clearly show distinct groups of individuals, the sex and life-environment repartition on the opposite, display equal variance between them (Levene’s test: PC1: F(3, 44) = 0.2445, p = 0.86; PC2: F(3, 44) = 0.3943, p = 0.76; PC3: F(3, 44) = 0.8425, p = 0.48; detailed in **Supplementary Table 1**). In conclusion, social profiles defined by unbiased clustering methods explain better the variance in social behaviours than sex and environment factors.

#### Definition of three distinct adolescent social profiles

To characterize each variable’s differences between the three groups, we used the variables from the PCA and performed statistical tests to compare the 3 groups for each social parameter (**Figure 2** & **Supplementary Figure 3**). Animals from group 1 was significantly expressing more dominance (KW(2) = 26.11, p < 0.0001) with a high number of approaches (KW(2) = 22.58, p < 0.0001) compared to the two other groups (Dunn’s test: group 2: dominance: p = 0.001; approach: p < 0.0001; group 3: dominance: p < 0.0001; approach: p = 0.0004). We thus named this group the dominant group. This group also significantly expressed less tolerance (KW(2) = 22.58, p < 0.0001), than the two other groups (Dunn’s test: group 2: p < 0.0001 ; group 3: p = 0.0004) and less interruption of contact (test mouse) (KW(2) = 8.367, p = 0.0152 ; Dunn’s test: group 2: p = 0.0467 ; group 3: p = 0.0084).Animals from group 2 on the other hand, significantly expressed more submission (KW(2) = 27.96, p < 0.0001) compared to the two other groups (Dunn’s test: group1: p < 0.0001 ; group 3: p = 0.0002). We thus named this group the submissive group. This group also significantly expressed higher motivation time (KW(2) = 8.4380, p = 0.0147) compared to the group 3 (Dunn’s test: p = 0.0365) and higher social discrimination (KW(2) = 8.9655, p = 0.0113) compared to the two other groups (Dunn’s test: group 1: p = 0.0150 ; group 3: p = 0.0180). Animals from group 3 only significantly expressed less contact time (KW(2) = 20.57, p < 0.0001) compared to the two other groups (Dunn’s test: group 1: p < 0.0001 ; group 3: p = 0.0013)(**Supplementary Table 1**).

Animals distinguished by their social behaviours also significantly differed for other variables not used in the PCA analysis, such as acoustic communication, paw control (from either animal) and for *exploration of the environment* (*i.e*., rearing, jumping, digging variables, **Supplementary Figure 3**). Animals of the dominant profile expressed significantly more paw controls (KW(2) = 12.79, p = 0.0017) compared to the two other groups (Dunn’s test: submissive: p = 0.0071; group 3: p = 0.0012). The dominant individuals also expressed enhanced communication (KW(2) = 9.321, p = 0.0095) compared to both other groups (Dunn’s test: submissive: p = 0.0152 ; group 3: p = 0.0091).

The submissive individuals, on the other hand, expressed more *submissive behaviour* (*i.e.* paw control from the new mouse)(KW(2) = 10.60, p = 0.005) compared to the two other groups (Dunn’s test: dominant: p = 0.0051 ; group 3: p = 0.0078).

Individuals from the group 3, not yet clearly characterised socially, were significantly less interested in social experience, but expressed increased *exploratory or anxiety-like behaviours* (jumping: KW(2) = 8.433, p = 0.0147; rearing: KW(2) = 10.03, p = 0.0066; digging (from the new mouse): KW(2) = 7.990, p = 0.0184), compared to dominant individuals (Dunn’s test: respectively p = 0.0074; p = 0.0025; p = 0.0075). We thus named this group the explorer group. These results highlight that animals from the explorer profile expressed more motivation, or at least the same level of motivation, for environment exploration than for social novelty.

The three profiles also significantly differed in other non social tasks, namely for *reward sensitivity* (sucrose preference) and *defensive reaction* (looming task), but not for locomotion (**Supplementary Figure 4 A, B & C**, **Supplementary Table 1**). Explorer individuals showed increased sucrose preference (Day1: KW(2) = 8.957, p = 0.0113; Day2: KW(2) = 9.220, p = 0.0099; Day3: KW(2) = 6.327, p = 0.0422) compared to dominant individuals (Dunn’s test: Day1, p = 0.0046; Day2, p = 0.0087; Day3, p = 0.0240) and submissive mice (Dunn’s test: Day2, p = 0.0177)(**Supplementary Figure 4 B**). Interestingly, explorer animals also explored significantly more during the habituation period before the looming task (middle: KW(2) = 6.880, p = 0.0321; opposite: KW(2) = 7.965, p = 0.0186) compared to the dominants (Dunn’s test: middle: p = 0.0212; opposite: p = 0.0076) that were more hiding than exploring (KW(2) = 6.806, p = 0.0333; Dunn’s test: p = 0.0156) (**Supplementary Figure 4 C**). These results together tend to support the hypothesis that explorer individuals might be more sensitive to their environment (i.e. exploration and sucrose reward) than their littermate peers in adolescence.

#### Social profiles: sex and life-environment distribution

In our cohort of animals 39.58% of individuals were explorers, 31.25% were dominants and 29.16% were submissives (**Figure 2 B**). This distribution, which was not completely equal, would tend to suggest that adolescents seemed more inclined to explore their new environment (*i.e.* 39.58% of explorers) than to interact with their new peer (*i.e.* 31.25% of dominants and 29.16% of submissive). For sex and life-environment repartition (**Figure 2 D**), most of the dominant individuals were females (80%) and from an enriched life-environment (60%). The submissive group was constituted predominantly by males (64.3%) and by mice raised in an enriched environment (71.4%). The explorer group was composed mainly of males (63.1%) and of mice from a standard environment (Sex: Chi² = 51.81(2), p < 0.0001; Environment: Chi² = 44.26(2), p < 0.0001; detailed in **Supplementary Table 1**).

### Adolescent social behaviours were associated with adult behaviours

Another objective was to investigate further if these adolescent profiles could be associated with adult behaviours. To answer that question, linear regression analysis was used from adolescent PCA scores to see association with adult social behaviours (**Figure 3** and **Supplementary Table 4**).

**Figure 3.**
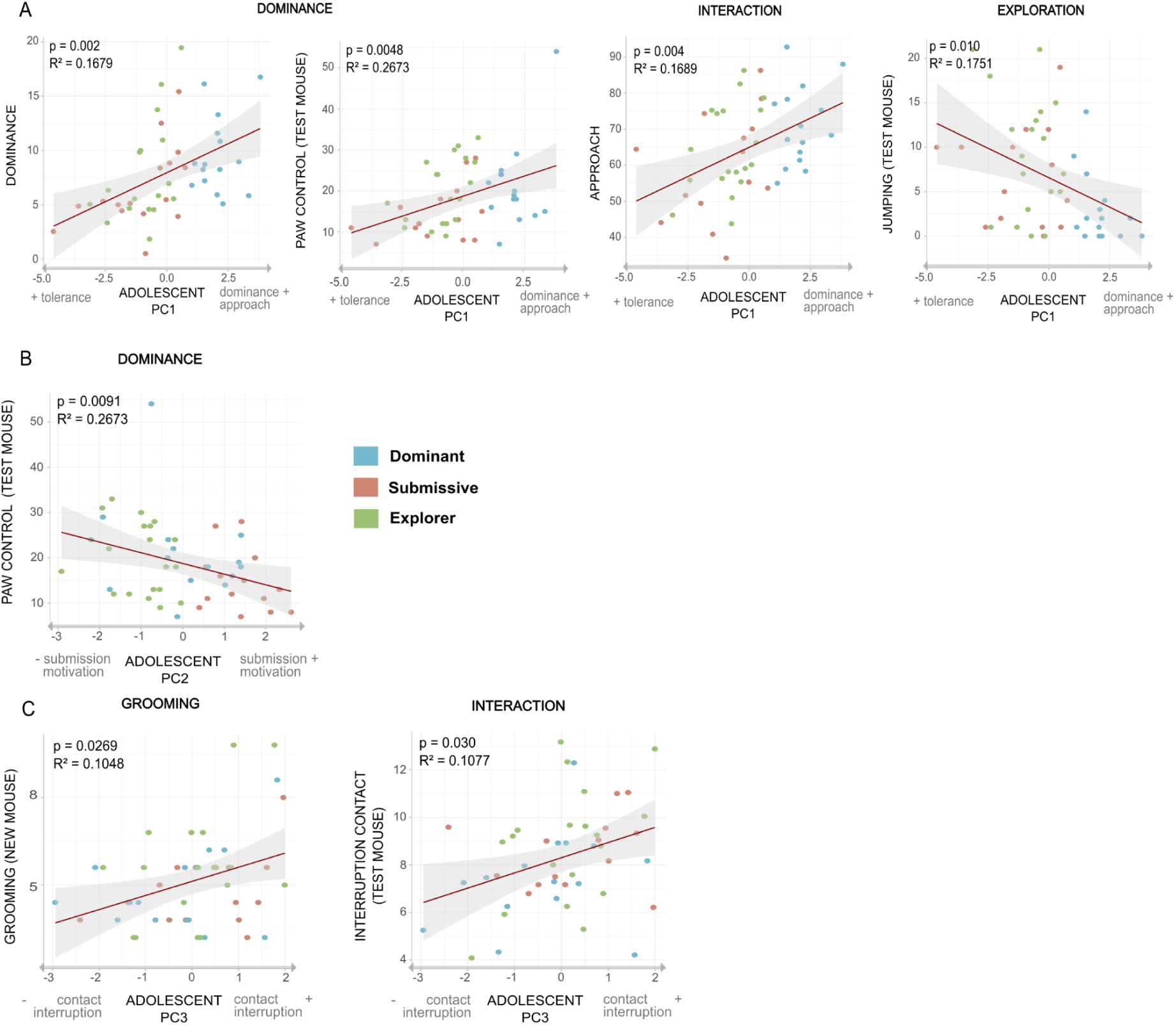
Adolescent social behaviours were associated with adult behaviours. Linear regressions between adolescent PCA scores, **A.** PC1, **B.** PC2 and **C.** PC3, and adult social behaviours for all social variables used in PCA: dominance (follow from test mouse and paw controls), interaction (approach and interruption of contact from test mouse), grooming (of the new mouse) and exploration (jumping from the test mouse). Each social profile is represented in colour. All significant and non-significant results are detailed in **Supplementary Figure 4**. Significative threshold after correction p < or equal 0.05.

The linear regression analyses showed that adolescent PC1 scores was positively correlated with adult *dominant behaviours* (dominance: R² = 0.1679, p = 0.002; paw control: R² = 0.2673, p = 0.005; **Figure 3 A**) and *interaction* (approach: R² = 0.1689, p = 0.004; **Figure 3 A**). Expectedly, it was also negatively correlated with tolerance (R² = 0.1689, p = 0.004; **Suppl. Figure 5 A**), *exploration or anxiety-like behaviour* (jumping : R² = 0.1751, p = 0.010; **Figure 3 A**) and rearing (R² = 0.1236, p = 0.021; **Suppl. Figure 5 B**), as well as *motivation* (motivation time: R² = 0.1629, p = 0.013; **Suppl. Figure 5 C**). For the adolescent PC2, which represents *submission* and *motivation behaviours*, it was negatively correlated with adult *dominant behaviour* (paw control: R² = 0.2673, p = 0.0091; **Figure 3 B**). Adolescent PC3 was positively correlated with interruption of contact in adults (R² = 0.1077, p = 0.03; **Figure 3 C**) and grooming from the new mouse (R² = 0.1048, p = 0.0269; **Figure 3 C**). Altogether, these results show that adolescent PCA dimensions previously used to establish the 3 social profiles are associated with adult social behaviours.

### Adult social behaviours were associated with specific gene expression patterns

The expression level of genes from neurotransmission systems known to be involved in social cognition was assessed in key brain structures, namely the prefrontal cortex, the cerebellum, the olfactory bulb, and striatal regions.

For the Olfactory bulbs, the individuals expressing more approach also significantly expressed less *Slc6a3* (DAT) (F(8, 39) = 1.95, p = 0.036, R²= 0.1391) and *Drd2* (F(8, 39) = 1.545, p = 0.032, R²= 0.085, **Figure 4**), and less *Drd1* (F(8, 39) = 2.491, p = 0.012, R²= 0.202), less *Bdnf* (F(8, 39) = 2.181, p = 0.024, R²= 0.167), less *Ngf* (F(8, 39) = 1.488, p = 0.011, R²= 0.077) and less *Ccnd1* genes (F(8, 39) = 1.282, p = 0.047, R² = 0.046, **Suppl. Fig. 6**), compared to individuals doing less approach. Similarly, other behaviours were predicting adult gene expression in the OB: the more dominant individuals expressed more *Slc6a4* (5-HTT) (F(8, 39) = 2.165, p = 0.017, R²= 0.166), more *Drd1* (F(8, 39) = 2.491, p = 0.0060, R²= 0.2024), more *Bdnf* (F(8, 39) = 2.181, p = 0.0189, R²= 0.1674) and more *Oxtr* (F(8, 39) = 1.408, p = 0.0400, R²= 0.0649) genes expression levels, compared to most submissive individuals (**Suppl. Fig. 6**). Interestingly, the dominance behaviours (dominance, approach) are predicting the OB gene expression levels, which could be explained by the fact that dominance is characterised by taking more information from the environment. There seems to be distinct differences between dominance (following the new mouse) and the approach (initiating contact with the new individual). More specifically, this result means that individuals that express more dominance (follow) also express more genes related to oxytocin receptor (*Oxtr*), monoamine receptors (*Drd1*) and transporter (*Slc6a4,* 5HTT) and growth factor (*Bdnf*), while individuals that express more approach (initiating contact) express less growth factor (*Ngf*, *Bdnf*) and less monoaminergic receptor (*Drd2* & *Drd1*) and transporter (*Slc6a3*, DAT).

**Figure 4.**
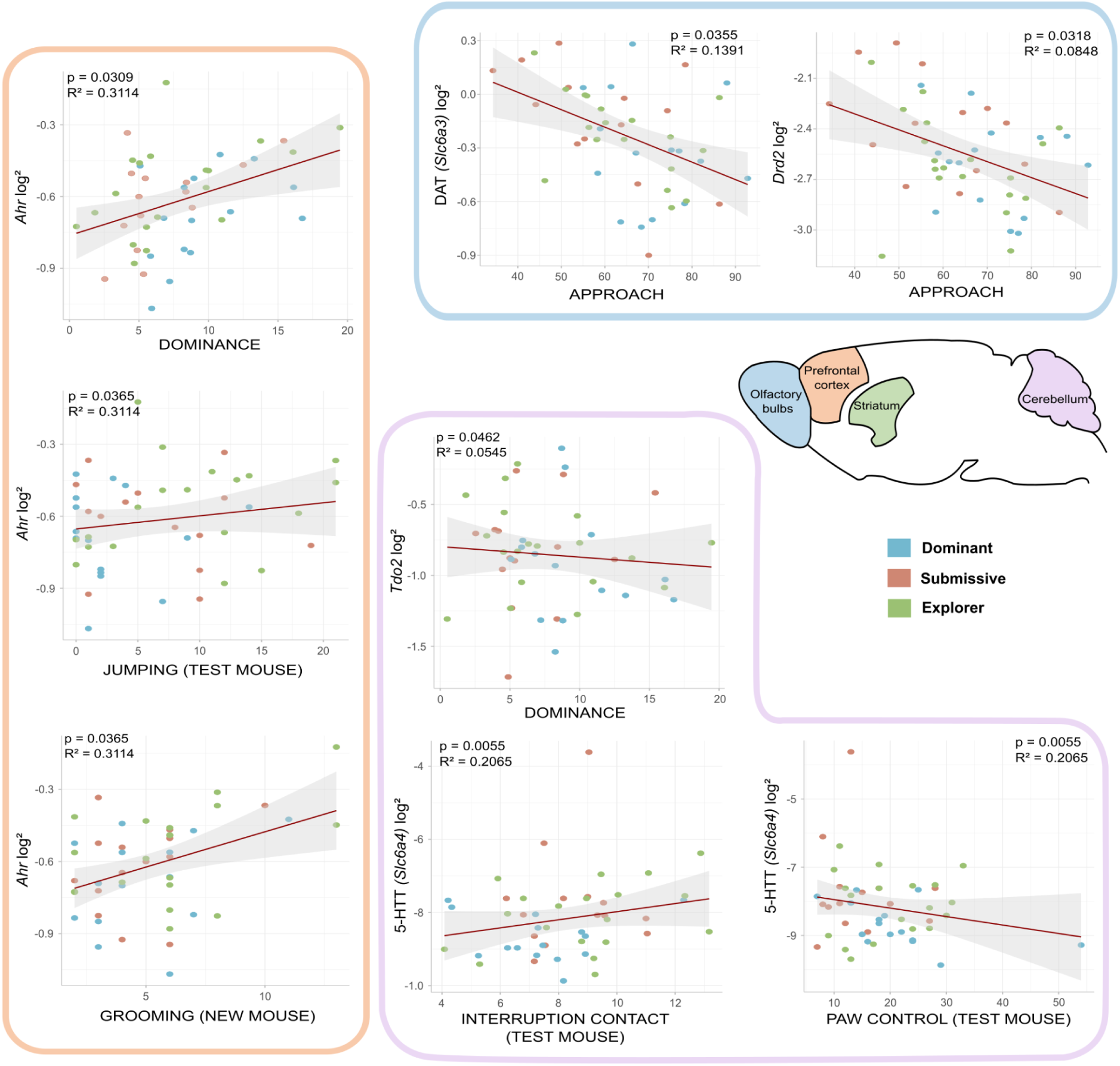
Adult social behaviours were associated with basal brain gene expression. Linear regression models from social behaviours at adulthood of RT qPCR levels in the brain. Analyses showed significant results for 5 genes in the Olfactory bulbs (in blue), the Prefrontal cortex (in orange) and the Cerebellum (in purple). All statistical results are reported in **Supplementary Figure 6** and in **Supplementary Table 5**.

In the Prefrontal cortex, mainly exploration behaviours are predictive of adult gene expression. For instance, the level of jumping from the test mouse significantly predicted *Ahr* (F(8, 39) = 3.657, p = 0.036, R²= 0.311, **Figure 4**) and *Avpr1a* (F(8, 39) = 1.644, p = 0.044, R²= 0.099, **Suppl. Fig. 6**) genes expression levels. In other words, individuals that do more exploration during interaction (i.e. jumping) will express significantly more *Avpr1a and Ahr* genes, while less exploring individuals will express it less. In addition, individuals that expressed more self-centered behaviours, such as grooming (from the new mouse) during interaction, also significantly expressed more *Ahr* gene (F(8, 39) = 3.657, p = 0.036, R²= 0.311; **Figure 4**). Finally, the most dominant individuals expressed significantly more *Ahr* gene (F(8, 39) = 3.657, p = 0.0309, R²= 0.3114; **Figure 4**).

For the Cerebellum, most dominant individuals (i.e. dominance, paw control) expressed significantly less *Tdo2* (F(8, 39) = 1.339, p = 0.046, R²= 0.054) and *Slc6a4* (5-HTT) (F(8, 39) = 2.529, p = 0.005, R²= 0.206; **Figure 4**). Individuals interrupting the contact more significantly expressed more *Slc6a4* (5-HTT) (F(8, 39) = 2.529, p = 0.005, R²= 0.206, **Figure 4**) and more *Ngf* genes (F(8, 39) = 3.037, p = 0.0498, R²= 0.257; **Suppl. Fig. 6**).. These results together suggest that most dominant individuals express less indoleamine pathway (*Tdo2*), monoamine transporter (*Slc6a4*, 5HTT) and nerve growth factor (*Ngf*) while individuals interrupting more the contact express more 5HTT (*Slc6a4*).

Concerning the Striatum, individuals expressing more approach (F(8, 37) = 4.289, p = 0.038, R² = 0.369) and interrupting the contact more (F(8, 37) = 4.289, p = 0.027, R²= 0.369) significantly expressed more *Htr2c* gene in opposition to most submissive individuals (**Suppl. Fig. 6**). Individuals that only expressed more approach also significantly expressed more *Oxtr gene* (F(8, 37) = 2.573, p = 0.038, R² = 0.219). For exploratory behaviours, individuals expressing more rearings significantly expressed more *Ido1* gene (F(8, 37) = 2.931, p = 0.009, R² = 0.256), while individuals doing more jumping during interaction, expressed less *Htr2c* gene (F(8, 37) = 4.289, p = 0.027, R²= 0.369, **Suppl. Fig. 6**). Also, individuals expressing more motivation time, significantly expressed more *Cdk5* gene (F(8, 37) = 2.362, p = 0.037, R²= 0.195), but less *Htr2c* gene (F(8, 37) = 4.289, p = 0.038, R²= 0.369) compared to submissive ones (**Suppl. Fig. 6**). These results show a certain prediction from dominance, interaction, motivation and exploration behaviours during interaction in adulthood of adult genes expression levels in different social brain areas. Overall, this suggests an impact for some individuals of adult social experience on general brain activity and the activity of specific signalling pathways.

As we showed that adult behaviors were associated with specific gene expression patterns, we assessed whether differences in gene expression levels were associated with differences in adult brain metabolites.

In the Olfactory bulbs, the level of interruption of contact significantly predicted the ratio of 5HTP/5HT and NA/Nor levels (**Suppl. Fig. 7**). More precisely, individuals that interrupted the contact more had more 5HT available, compared to individuals that interrupted the contact less which had more 5HTP available (F(8, 38) = 2.17, p = 0.014, R²= 0.169, **Suppl. Fig. 7**). At the same time, individuals that interrupted the contact more also had a higher availability of Noradrenaline in contrast to individuals interrupting the contact less which had Normetanephrine more available (F(8, 38) = 1.63, p = 0.032, R²= 0.099, **Suppl. Fig. 7**).

Finally in the PFC, individuals that interrupted the contact more during interaction had more 5HT available compared to individuals that interrupted the contact less which had 5HTP more available (F(8, 31) = 1.9, p = 0.037, R²= 0.155, **Suppl. Fig. 7**).

Interestingly, the level of adult brain metabolites is only predicted from the level of interruption of contact during adult interaction. These results show interrupting social contact at adulthood is significantly associated with brain metabolites levels in specific brain areas belonging to the social brain, suggesting an impact of adult social experience on overall brain activity and the activity of specific signalling pathways. However, we cannot rule out the fact that brain regional metabolite contents were structured and shaped at adolescence, thus leading to adult social behaviours. Our last objective was to assess if adolescent social behaviours could already be associated with neurobiological variations, and whether from them we could predict the level of some adult neurobiological features.

### Adult brain gene expression patterns are associated with adolescent social behaviour

As adult social behaviour can predict adult neurobiological features we wondered whether some specific adult behavior and/or biological features could already be associated with adolescent behaviour (**Figure 5**, **Supplementary Table 7**).

**Figure 5.**
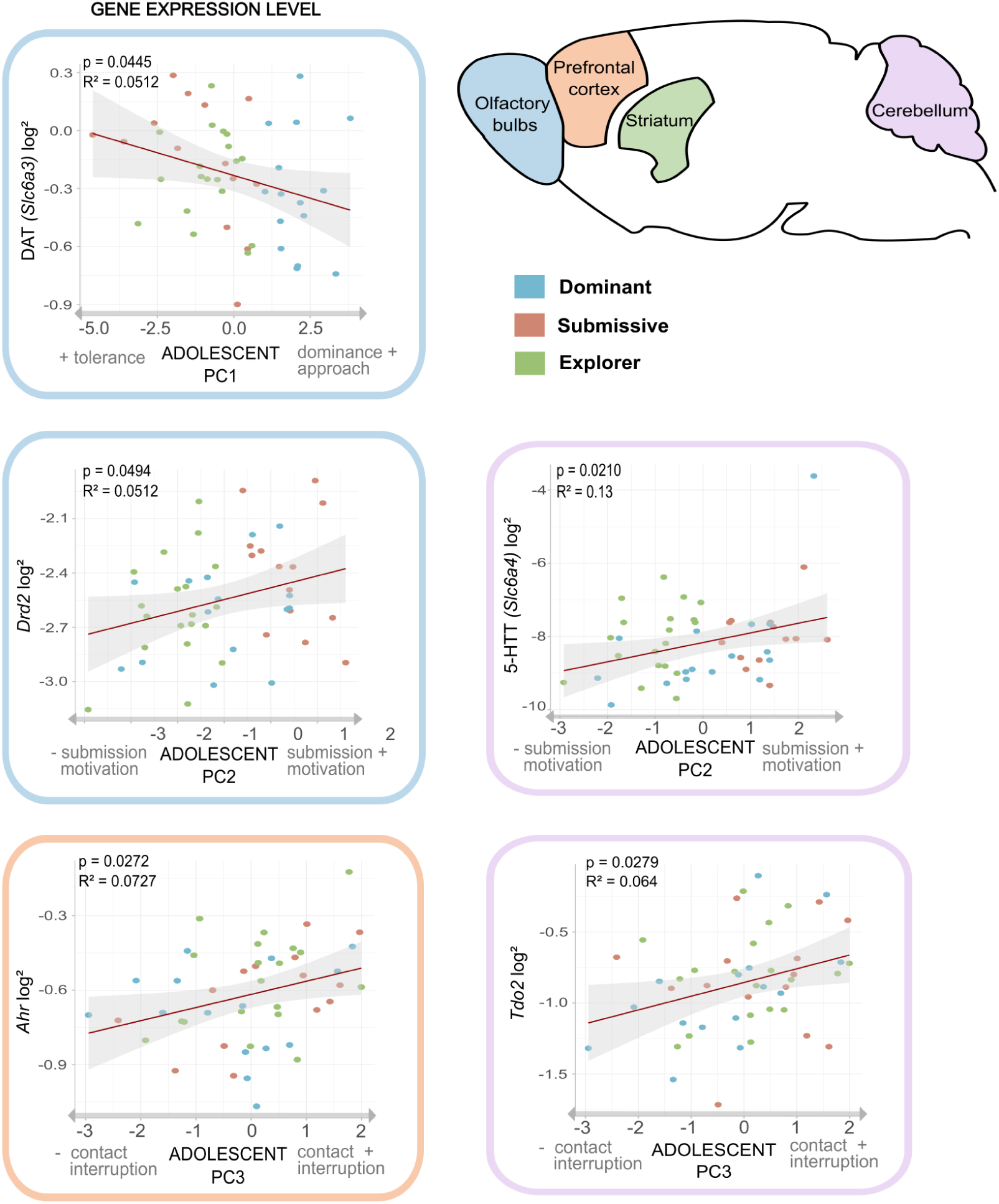
Association from adolescent social profiles with adult brain features. Linear regression models of adolescent social PCA scores and brain regional RT qPCR levels. Analyses showed significant association with the expression of 5 genes in the Olfactory bulbs (blue), the Prefrontal cortex (orange) and the Cerebellum (purple). All results are detailed in **Supplementary Table 7** along with other results that didn’t reach statistical threshold (p⩽0.05).

The level of adult gene expression was associated with adolescent PCA scores -based on social behaviours (**Figure 5**). Adolescent mice that were more likely to interrupt social contacts (PC3, i.e., interruption of contact during the social task) exhibited a higher activity of the indoleamine pathway (*Ahr*) in the PFC (F(3, 44) = 2.23, p = 0.0272, R²= 0.0727) and more kynurenine synthesis gene (*Tdo2*) in the cerebellum (F(3, 44) = 2.07, p = 0.0279, R²= 0.064).

By contrast, adolescent mice that were more submissive and motivated for social contact (PC2, i.e., submission and motivation in social tasks) were more likely to express a higher level of *Drd2 in the* olfactory bulbs when adult (F(3, 44) = 1.84, p = 0.0494, R²= 0.0512) and of *Slc6a4* (5HTT) gene expression in the cerebellum (F(3, 44) = 3.35, p = 0.0210, R²= 0.13). Also, mice that were more dominant and interacting at adolescence (PC1, i.e., dominance and interaction) were less likely to express *Slc6a3* (DAT) gene expression when adult compared to mice with low dominance level (F(3, 44) = 1.84, p = 0.0445, R²= 0.0512).

Overall these results suggest an impact of early social profiles on regional brain activity, particularly in brain areas devoted to social behaviors, potentially influencing (or reinforcing) the development of specific signalling pathways.

## Discussion

We propose that it is possible to unveil social profiles at adolescence that predict social behavior at adulthood in association with regional gene expression and monoamine content. Using PCA analysis on multiple social variables we determined three social profiles at adolescence: (1) a dominant profile expressing significantly more dominance and acoustic communication, (2) a submissive profile expressing more submissive behaviour and social motivation, (3) an explorer profile neither motivated nor interested in social interaction. The three profiles also differ regarding non-social behaviors such as novelty exploration level and reward sensitivity.

Animals with the dominant profile show more dominant parameters, communication, a parameter that we previously showed to be associated with dominance parameters (Chabout et al., 2012; Coura et al., 2013; Simola & Granon 2019, for review). Dominant mice during social tasks also show decreased exploration of all novel environments and spend more time hiding when possible, suggesting more anxiety-like behavior. They also consume less sucrose than other groups, particularly during the first two days of exposure to this novel taste, suggesting a tendency to neophobia.

It is interesting to note that animals with a dominant profile are mainly females (80%, or 46.66% enriched and 33.33% standard) that are more neophobic than male (Greiner & Petrovich 2020) suggesting that neophobia may be a major constituent of social behavior, particularly when considering the establishment of a novel social interaction with an unknown conspecific. We can’t also ignore that this distribution might be attributed to dominance expression within same sex dyads could differ between males and females. Since female mammals are also reported to be more sensitive to olfactory cues (Baum & Keverne, 2002; Kass et al., 2017) and that hierarchy information in rodents is mainly conveyed by urine olfactory signals (Borak et al., 2025), we chose to focus on olfactory brain areas to measure gene expression and monoamine markers that will be discussed below. Animals with a submissive profile were mostly constituted of males (35.7% females and 64.28% males). They were more motivated by the discovery of a novel conspecific and tolerate significantly more being approached by an unknown congener.

Animals showing the explorer profile favor discovering novel environments over novel social partners, suggesting that they may be more sensitive to their environment (i.e. exploration and food reward) than their peers in adolescence (Adcock et al., 2021). Seventy percent of explorer animals were reared in a standard environment -although not all animals reared in standard cages belong to explorer profile. It is thus plausible that living in a standard (rather ‘impoverished’ here) environment, increases interest in novelty exploration (Lin et al., 2011; Mesa-Gresa et al., 2013) because of the sensory stimulation it provides. The results show that sex and environment factors did not explain the variance of the data, while social profiles did, suggesting that, although rearing environment and sex factors may influence social profiles, they are not sufficient. From the repartition of individuals between these three profiles, we could suggest that adolescent mice seem to be more motivated by novelty exploration (*i.e.* 39.58% of explorers) rather than interacting with novel same sex congeners, likely because at adolescence, mice live in groups and have never been socially isolated before social tasks. Indeed, we previously showed that social isolation before social tasks promote social contact (Avale et al., 2011) and acoustic communication (Chabout et al., 2012; Lefèbvre et al., 2020).

Social variables linked to adolescent profiles were also associated with social behavior of the same animal at adulthood. This is particularly true for the level of dominance and social interaction and motivation, novelty exploration and self-centered behavior such as self grooming. Our data suggest that dominance tendencies already present at adolescence are likely to stabilize through lifespan, thus influencing and shaping future social behaviors. Individual stability in social behaviour such as hierarchical dominance and interaction, is important at group level for evolutionary consideration, as it promotes individual fitness, better feeding and mating opportunities, access to reinforcing stimuli and has some health benefits (McNamara et al., 2018, for more detailed review). On the contrary, social and dominance instability (resulting in partner changes) has been shown to promote anxiety, stress and reduced breeding (McNamara et al., 2018 for review). The ability of a mouse to show dominance and to develop peaceful interaction with a novel conspecific is part of social abilities used to develop hierarchy in a group. The fact that this ability develops from adolescence and is mainly stable within individuals also stabilises relationships between individuals in social groups, leading to global fitness of the group. Individual stability of social features from adolescence to adulthood led us to wonder whether this stability is supported by individual neurobiological and/or gene expression features.

In adult animals high level of dominance was associated with higher gene expression in the olfactory bulb (OB) of oxytocin receptor (*Oxtr*), of monoamine receptors (*Drd1*), of 5-HT transporter (5HTT) and of growth factor (*Bdnf*). By contrast, approach behavior was correlated with lower expression of growth factor (*Ngf*, *Bdnf*) monoaminergic receptor (*Drd2* & *Drd1*) and dopamine transporter (DAT).

OB plays a crucial role in the initial complex social information processing (Lazzarini et al., 2014), and is also richly innervated in dopaminergic neurons (Pignatelli & Belluzzi, 2017). These dopaminergic neurons do not seem to be required for essential olfactory processing (Akarawa, 2021) but rather implicated in social olfactory discrimination. Activation of D2 DAR, or lack of dopamine transporter DAT induce odor discrimination deficit compared to wild type animals (Tillerson et al., 2006, Escanilla et al., 2009). Lower expression of *Drd2* and DAT genes lead to greater dopamine level and neuronal excitability in the olfactory bulb. Similar effect is found in DAT KO mice where absence of DAT expression leads to 5 time increase of DA and downregulation of D2R in striatum (Giros et al 1996). This increased Dopamine and reduced D2R expression might increase D1R activation by dopamine and thus be beneficial for odor discrimination (Escanilla et al., 2009). D1 receptor activation might have opposite effect on odor discrimination, as pharmacological studies demonstrate opposite effect of D2 or D1 agonists (Doty et al., 1998, Yue et al., 2004) and that D1 activation of glomerular circuit elevates the system’s sensitivity to odor stimuli (Liu, 2020). Interestingly, as oxytocin increases social interaction and recognition through effects on olfactory bulb, we might expect elevation of oxytocin receptor expression to increase olfactory ability and social interaction in dominant animals (Sun et al, 2021, Grinevich et al., 2018). An hypothesis might be that this improved odor discrimination gives an advantage in the understanding of social cues, leading in turn to more appropriate social interaction (in our case, dominant behavior as our social task promotes dominance). Indeed, integration of olfactory cues has been shown to be key for the development of social behavior particularly at early ages when other senses are less mature, in humans and other animals (for review Damon et al., 2021; Ervin et al., 2015; Keller-Costa et al., 2015).

Approach and dominant behaviour predict lower expression of indoleamine pathway enzyme (*Tdo2*), monoamine transporter (5HTT) and nerve growth factor (*Ngf*) in the Cerebellum. Tryptophan could either enter in the pathway of 5-HT synthesis or in the kynurenine pathway. With IDO1, TDO2 is another limiting enzyme that catalyzes the first step of the kynurenine pathway, especially in the cerebellum (Mazarei et al., 2013; Yuasa et al., 2009) . Even if the majority of tryptophan enters the kynurenine pathway, a lower expression of *Tdo2* gene that might induce a lower activation of the kynurenine pathway potentially increases the availability of tryptophan for 5-HT synthesis. Moreover, In the brain, 5HTT is a protein allowing reuptake of 5HT from the synaptic cleft into serotonergic neurons, thus having a central role in fine-tuning serotonin neurotransmission. Low amounts of 5HTT are thus expected to lead to higher synaptic and extracellular 5HT levels. Our results thus support a relationship between cerebellar 5-HT regulation and dominant behavior.

Approach and dominant behaviour also predict greater expression of (Ahr) in the Prefrontal cortex. Ahr (aryl hydrocarbon receptor) is a transcription factor quite known to be critical for organismal homeostatic functions, as it senses environmental and endogenous compounds, and initiates genes expression programs influencing metabolism, development, inflammation and immune response (Anderson et al., 2013). Polymorphisms of Ahr-related genes are related with the severity of autistic symptoms especially for social communication (Fujisawa et al., 2016). Further research will be needed to decipher how Ahr modulation of gene expression programs in the prefrontal cortex might predict social behavior.

Our results show that the more dominant mice are at adolescence, the more dominant and approaching they will be at adulthood, and the less they will express dopamine transporter (*DAT)* and Drd2 in the olfactory bulb. The more mice interrupt social contact at adolescence, the more they interrupt contact and perform self-grooming at adulthood and activate the kynurenine pathway (*Ahr gene expression*) in the Prefrontal cortex and indoleamine synthesis gene (Tdo2) in the Cerebellum. Interestingly, the less mice show social motivation at adolescence, the less they will be dominant at adulthood, and the more they will express the monoamine transporter (5HTT) in the cerebellum.

One question might be to understand why having this behavior at adolescence might predict such gene expression. Our data show that being more dominant, interacting and socially motivated might shape olfactory activity at adolescence, whereas animals more interested in novelty exploration or socially submitted would develop other abilities. Interestingly, DA interneurons of OB are highly plastic in relation to activity and are mostly continuously renewed by neurogenesis through life (Bonzano et al., 2016; Lazarini et al., 2014). For example, enrichment of the life environment, comprising frequent social interactions with multiple partners, downregulates DAT (*Slc6a3)* expression, that consequently leads to increase of synaptic occupancy of dopamine (Lee et al., 2013). In our results, most dominant adolescents express less *Slc6a3* gene at adulthood, a biological feature that might be related to environmental enrichment from adolescence to adulthood. The fact that 60% of animals with a dominant profile were reared in an enriched environment suggests that one of the factors that may influence or reinforce the regulation of the dopaminergic system in the olfactory bulbs is the opportunity to have a richer social life. Alternatively, animals with this social profile may have a pre-existing low DAT and/or low Drd2 gene expression. Therefore, we propose that adolescent individuals as adolescents that express low DAT and *Drd2,* which would lead to an increased release and use of DA in the OB, are more likely to be more prone to interact with new conspecifics. By contrast, animals that are submissive as adolescents that express more DAT and more *Drd2*, which would lead to less release and use of dopamine in the OB, would be less prone to explore novel conspecifics.

In humans an association between 5HTT and brain areas volume has been shown, which underlies individual differences in emotion processing and constitute a vulnerability feature to emotional disorders (Radua et al., 2014). Also, individuals with COMT (catecholamine transferase, degradation monoamines) and 5HTT specific allele polymorphisms association showed smaller gray matter volume in the cerebellum and other structures, taken as indicators of emotional disorders and a vulnerability trait to major depressive disorders (Kempton et al., 2011). In addition, genetic variation of 5HTT was shown to be at the onset of individual variability in the modulation of specific brain circuits (amygdala, PFC and cerebellum) in response to negative emotional stimuli (Canli et al., 2005), and is linked to anxiety-related personality traits such as neuroticism and harm avoidance (Canli et al., 2005). This effect of 5HTT and COMT on individual variability and predisposition to pathological traits can be explained by the role monoamines play in synaptic and brain plasticity during cortical maturation (Radua et al., 2014). More specifically, dopaminergic innervations have been found to influence the course of cortical development (Rosenberg and Lewis, 1995) while serotonin influences the development of the cortex and hippocampus (Zhang et al., 2006; Choi et al., 2007).

Our data in mice support 5HTT gene expression in the cerebellum as a major actor of individual differences in emotion processing. All together, we could suggest that individuals that are more submissive although socially motivated as adolescence might perceive social stimuli less positive than other individuals, a social trait potentially associated with other anxiety-like behaviors.

In our results, we suggest that adolescent individuals interrupting more contact will express more Ahr in PFC and more Tdo2 in Cerebellum when adults. Tdo2 and IDO1 are two rate-limiting enzymes of the kynurenine pathway (Higuchi & Hayaishi, 1967; Yuasa et al., 2009). Interestingly, Tdo2 is especially expressed in the cerebellum (Mazarei et al., 2013). Interestingly, products of the kynurenine pathway like kynurenine or kynurenic acid, are known agonists of the Aryl hydrocarbon receptor (AhR) transcription factor. Ahr (aryl hydrocarbon receptor) is a transcription factor involved in metabolism, development, inflammation and immune response (Anderson et al., 2013). If we consider that having more interruption of contact means a more stressful and negative social contact, thus we might suggest a possible link between these negative social experiences at adolescence and kynurenine pathway in PFC and cerebellum once adult.

## Conclusion

In conclusion, our results suggest that behavioral factors that explain better the variance of social data, hence delimitate social profiles are related to olfactory abilities and quality of social contact rather than to sex and life environment. Accordingly, we suggest that the typologies of social experiences in adolescence, either experienced in a positive way by rather dominant individuals, or experienced in a rather negative way by rather submissive individuals, predict social characteristics in adulthood. These adolescent social experiences might either shape activity of the dopamine system within olfactory bulbs and thus olfactory abilities, and/or efficiency of the kynurenine pathway in the PFC-Cerebellum network. Moreover, some genetic variability, like 5HTT expression in the cerebellum, might be already present at adolescence sustaining social behavior and profiles.

## Material & Methods

### 2.1. Animals

48 C57BL/6J mice (from Charles River, Lyon, France) were used for behavioural tests, measurement of brain monoamines (HPLC), and microfluidic RT qPCR. For this work, we studied 24 males and 24 females, 12 males and 12 females grown in enriched environments, and 12 males and 12 females maintained in standard environments (see below 2.2 for details). These animals were followed and tested throughout their development, from adolescence (34 to 57 days) to adulthood (78 to 93 days). The procedure and environments are detailed in **Figure 6**. No animal was deprived of any sort: of food, water or socially isolated prior to any experimental task.

**Figure 6.**
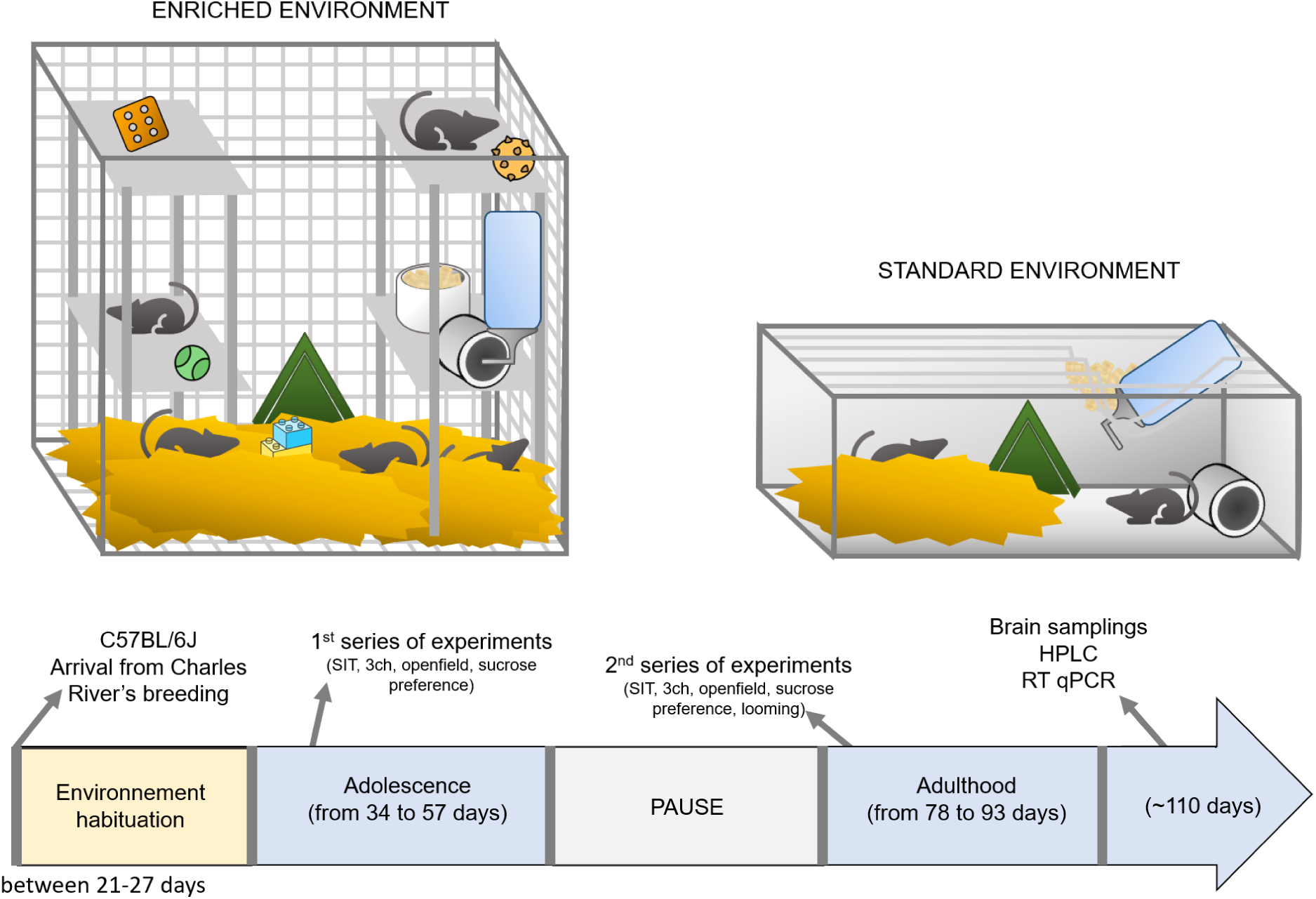
Representation of enriched and standard life-environment and a graphical representation of the procedure time frame.

Animals were treated according to the ethical standards defined by the Centre National de la Recherche Scientifique for animal health and care with strict compliance with the EEC recommendations (APAFIS n°4514-2021121622122648 v6).

### 2.2. Procedure

#### Type of environment: enriched vs. standard

We focused on the influence of the environment of life -rich versus standard- on the development of the social repertoire and on variability detection. To do so, after weaning (between 21 and 27 days), mice were placed in their respective environments under a temperature of 20-22°C with a day-night cycle of 12/12h and a luminosity between 270/300 lux for standard conditions, or 100/125 lux for enriched conditions. For the standard life-environment, mice were in groups of 4 individuals of same gender and age. The 15×30 cm cages had a ‘poor’ enrichment (**Figure 6**) with a small house, a cardboard tunnel and cotton. The ‘new’ mice that served as social stimuli during the social motivation and social interaction tasks came exclusively from this type of environment. Whereas, for the enriched environment, mice were in groups of 12 individuals of the same gender and age. The 74×45×72 cm cages were enriched with ‘rich’ objects, such as toys, objects (sound, textured or mobile), boxes, tunnels and different smells, which were changed twice a week to maintain novelty.

### 2.3. Behavioural Apparatus & Experimental Protocols

#### Three-chambers task (3-ch)

**Apparatus:** as described before in Lhopitallier and coll. (2022), we used a rectangular transparent Plexiglas cage divided into three equal compartments (64 cm X 42 cm). The apparatus contained two empty pots (diameter 9.2 cm; H 10.2 cm) placed at each opposite ends of the cage, with one of them containing a mouse (same sex and age as the test mouse hereafter called the “new mouse”) while the other one remained empty (see **Figure 7** in **Methods**). Cameras (Hercules®) were used to record the different behaviours of the ‘test’ mouse within the entire environment. Luminosity was set at 100 Lux and temperature at 20-22°C.

**Figure 7.**
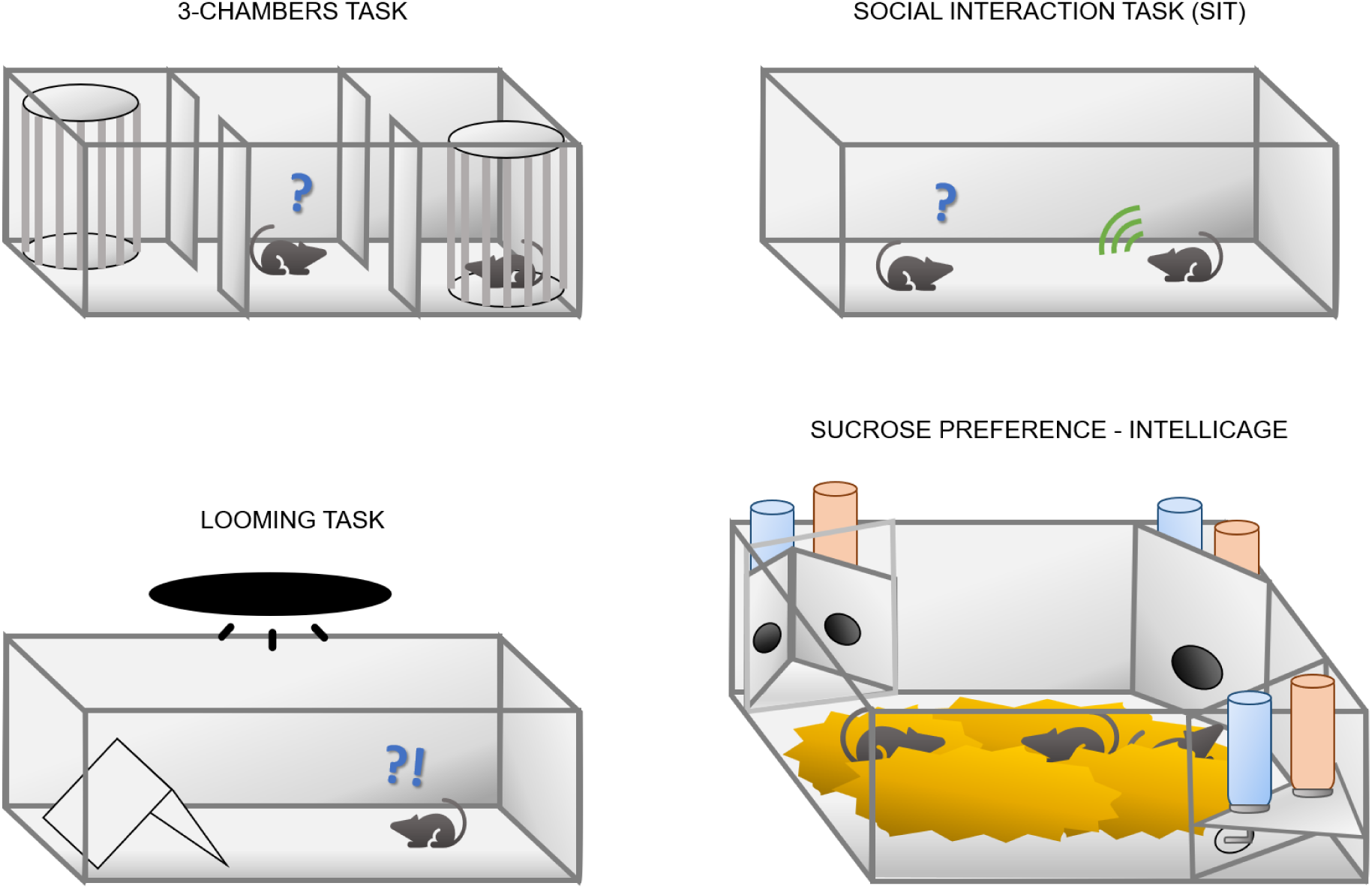
Representation of social tasks apparatus (social motivation 3-chambers task, social interaction task (SIT)), sucrose preference (lower right) and looming task (lower left).

**Protocol:** before starting the test all mice (‘test’ and ‘new’) were habituated to the experimental room for 15 minutes. Then, the “test” mouse was placed in the three-chamber apparatus for 15 minutes, allowing it to freely explore the environment and get used to the environment and the two empty pots. After this period of habituation, the ‘test’ mouse is kept in the central chamber by Plexiglas windows while the ‘new’ mouse is introduced into the ‘social jar’. The Plexiglas windows were removed so that the test mouse can roam around. Between each test mouse, the jars placement was counterbalanced and the odours homogenised by cleaning the apparatus with water to avoid any olfactory bias.

**Parameters measured:** we recorded behaviours during 10 min with ANY-maze software. The parameters analysed off-line concerned: total contact time with the ‘social jar’ for the ‘test’ mouse (sec) and the ‘social motivation’ ratio (contact time in the ‘social chamber’ (sec) / contact time in the ‘non social chamber’ (sec)). The time spent in contact with the jars was evaluated when the ‘test’ mouse is close enough or pokes with its nose the jar. All the social motivation parameters measured are described in **Table 8** (in **Methods**).

**Table 8.**
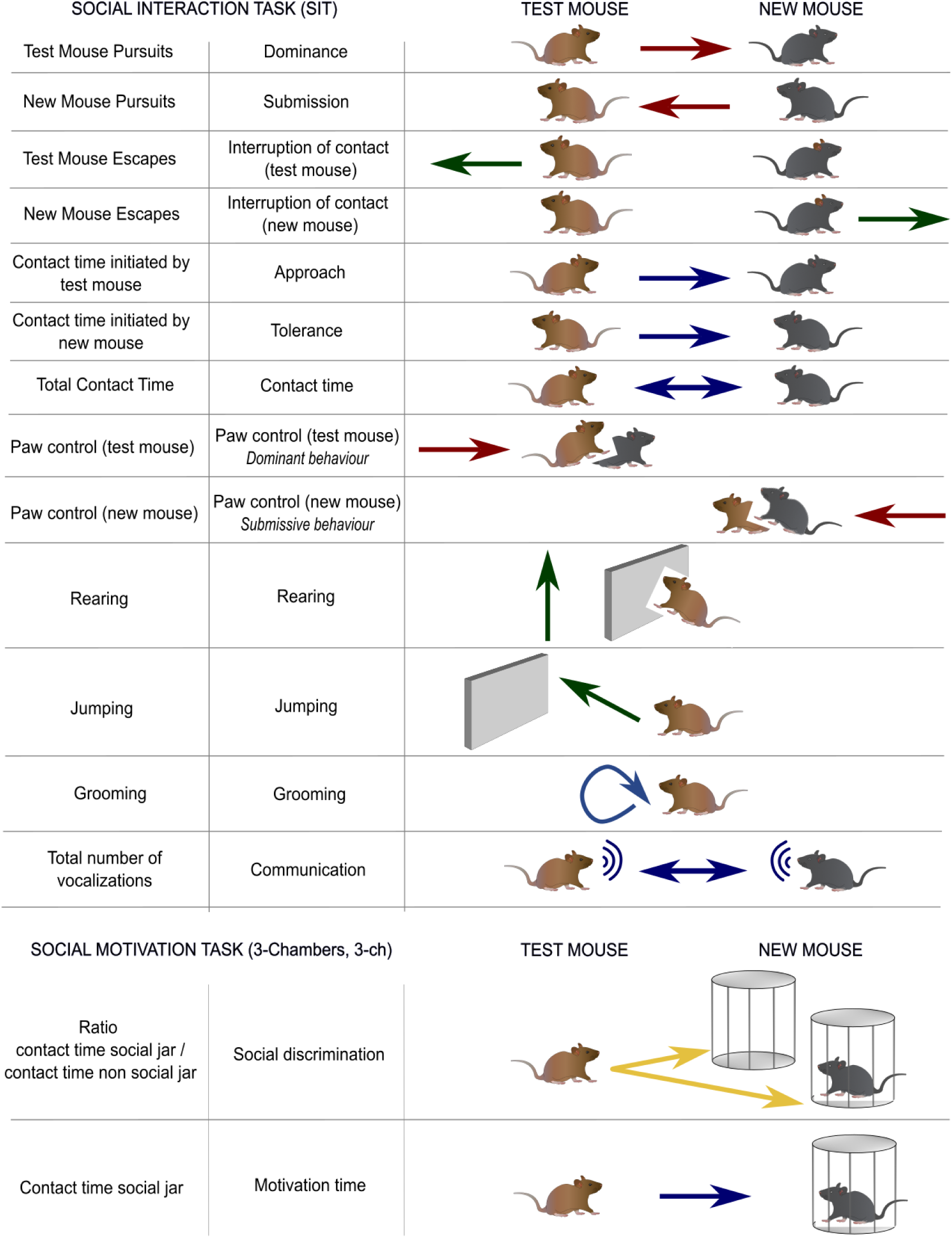
Summary table detailing all social behaviours measured for the Social Interaction Task (SIT) and the social motivation task (3-chambers task).

| SOCIAL INTERACTION TASK (SIT) |  | TEST MOUSE | NEW MOUSE |
| --- | --- | --- | --- |
| Test Mouse Pursuits | Dominance |  |  |
| New Mouse Pursuits | Submission |  |  |
| Test Mouse Escapes | Interruption of contact (test mouse) |  |  |
| New Mouse Escapes | Interruption of contact (new mouse) |  |  |
| Contact time initiated by test mouse | Approach |  |  |
| Contact time initiated by new mouse | Tolerance |  |  |
| Total Contact Time | Contact time |  |  |
| Paw control (test mouse) | Paw control (test mouse)<br>*Dominant behaviour* |  |  |
| Paw control (new mouse) | Paw control (new mouse)<br>*Submissive behaviour* |  |  |
| Rearing | Rearing |  |  |
| Jumping | Jumping |  |  |
| Grooming | Grooming |  |  |
| Total number of vocalizations | Communication |  |  |
| SOCIAL MOTIVATION TASK (3-Chambers, 3-ch) |  | TEST MOUSE | NEW MOUSE |
| Ratio<br>contact time social jar /<br>contact time non social jar | Social discrimination |  |  |
| Contact time social jar | Motivation time |  |  |

#### The Social interaction task (SIT)

**Apparatus:** for the Social Interaction Task we used a rectangular openfield made of transparent Plexiglas (50cm x 30cm x 30cm) that we described in details previously (e.g., Granon et al., 2003; de Chaumont et al., 2012; Nosjean et al., 2015; **Figure 7** in **Methods**). Luminosity was set at 100 Lux, temperature at 20-22°C and Cameras (Hercules®) were used to record the different behaviours.

**Protocol:** briefly, it contains a handful of clean bedding for each session and the ‘test’ mouse is placed in it for a 30-min habituation period before a same sex, same age unfamiliar conspecific (thereby called the ‘new’ mouse) is introduced in the openfield for a 8-min interaction. Prior to the test, the test and new mice got used to the experimental room for 15 minutes.

**Social parameters measured:** this task allows the assessment of social decision making when a mouse is exposed to a novel unfamiliar conspecific in a novel environment and thus is susceptible to make different kinds of choices (social versus exploration of novelty). We recorded behaviours during 8 min with video recordings using the Solomon coder ® software (solomoncoder.com) allowing real time and scalable ethological keyboard. The parameters analysed off-line concerned: social contact time (in sec, initiated either by the ‘test’ or ‘new’ mouse and the total contact time from both), following behaviours and interruption of contact (sec), dominant and aggressive behaviours (number of paw control, aggressiveness, tail rattling), exploratory activities (number of rearing) and anxiety-like behaviours (jumping, number of stop, digging, grooming), all those behaviours from both mice. In this task, we measure all parameters from the ‘test’ mouse point of view, and that of the ‘new’ mouse only in relation to the host mouse, which remains the reference mouse as it was the one we forced the dominance.

**Communication parameters measured:** during the SIT task, mice emitted ultrasonic vocalisations (USVs) that were recorded during the 8 min with a condenser ultrasound microphone Polaroid/CMPA placed above the apparatus (Lhopitaller et al., 2022). The microphone was connected to an ultrasound recording interface Ultrasound Gate 416H from Avisoft Bioacoustics H (Berlin, Germany), itself connected to a computer equipped with the recording software Avisoft RecorderUSG (Sampling frequency: 250 kHz; FFT-length: 1024 points; 16-bits). USVs were analysed off-line with SASLab Pro (Avisoft Bioacoustic H, Berlin, Germany). Spectrograms were generated for each detected call (Sampling frequency: 250 kHz; FFT-length: 1024 points; 16-bit; Blackman window; overlap: 87.5%; time resolution: 0.512 ms; frequency resolution: 244 Hz). All the social interaction parameters measured are described in **Table 8** (in **Methods**).

**Locomotion parameters measured:** during the 30-min habituation phase prior to the task, we analysed off-line the following parameters: total distance travelled (meters) and total mobility time (sec).

#### Defensive reaction to a threatening visual stimulus: Looming task

**Apparatus:** the objective of the looming task is to trigger and measure the reactivity of a mouse in response to a brief and sudden looming visual stimulus that mimicks a predator, which was previously described more thoroughly (Lhopitallier et al., 2022; Yilmaz et al., 2013). For that, we used a rectangular empty Plexiglas cage (L 50cm x 25cm x h 30cm) containing an opaque refuge in the extreme left (**Figure 7** in **Methods**). The cage was covered by a computer screen to display the visual looming stimulus directly above the animal at a diameter of 2 degrees of visual angle. Behaviours were recorded with a camera (Logitech) placed at 40 cm in front of the cage. During all the experiment, the diffuse light was set at 100 lux.

**Protocol:** we placed a mouse in the apparatus in order to let her get habituated to the arena for 10 min. Then, we triggered a visual stimulus when the mouse passed near the centre of the arena, which consists of a black disk rapidly increasing in size. The stimulus expanded to 20 degrees in 250 ms, and remained at that size for 250 ms. This stimulus was repeated 15 times and total exposure lasted 15 seconds (Yilmaz et al., 2013).

**Parameters measured:** we measured off-line: the time spent (in sec) in each section of the apparatus (under the shelter or ‘hiding’, ‘shelter’ area, ‘middle’ and at the ‘opposite’ bottom) during the habituation phase and post-looming stimulus phase, plus the reaction to the looming stimulus (in sec : freezing, hiding, tail rattling and reaction time or ‘RT’).

#### Food reward motivation: Sucrose preference test

**Apparatus:** The test was carried out in Intellicages homecages (TSE-System®, Bad Homburg, Germany), which consists of a rectangular Plexiglas transparent cage (55cm x 38cm x 21cm) (Iman et al., 2021). In each corner is placed a triangular box (15cm x 15cm x 21cm) which contains two side-doors each giving access to a bottle. Animals were housed in groups of 12 (one cage of ‘enriched’ mice and another of ‘standard’ life-environment; **Figure 6** in **Methods**) and were individually tracked because they were equipped with RFID chip detection systems. The Intellicages enabled us to monitor animals in their housing cage while they were performing the task, without interfering with the experiment.

**Protocol:** First, animals were habituated to the apparatus for 24 hours, and then the sucrose preference test ran for 3 days. During the habituation phase, there was only water in every bottle, then during the sucrose preference test, they had the choice in each corner between one bottle of water and one of sucrose 1% (1g/100mL). The order of the bottles was counterbalanced each day of the test.

**Parameters measured:** all behavioural parameters were monitored and analysed by the Intellicage Plus Software (“Analyzer”, TSE-System®) that enabled to assess: the number of licks of water and sucrose for each day, and the mean of water and sucrose consumption. We then assessed individual sucrose preference as an index for sensitivity to food reward (Pittaras et al., 2016; Ping et al. 2012). The sucrose preference score was thus calculated by: [(number of sucrose solution licks) / (number of sucrose solution licks + number of water licks)] × 100.

### 2.4. Statistical & Coding Analysis

#### Statistical analyses

Statistical analyses were carried out using the Rstudio software (version 4.2.2 of the R foundation for Statistical computing with Rcmdr-package).

Beforehand, each **variable distribution** was tested with the Shapiro-Wilk test to attest that most variables didn’t follow a Gaussian distribution (**Suppl. Table 1**). Thereafter they were tested with the non-parametric tests or normalised and corrected to carry out parametric analyses.

For **behavioural differences** between the social profiles, Kruskal-Wallis tests were used, and when results were significant, post-hoc analyses were conducted with Dunn’s multiple comparisons test.

The Levene’s test was used to test for homogeneity of variances between social profiles and sex and environment groups. And a Chi² analysis was performed to statistically check significant differences in the repartition (in %) between sexes (male vs. female) and environment (standard vs. enriched) for each social profile.

Concerning the **prediction analyses**, the behavioural variables were standardised and centred, and the gene expressions and basal metabolites levels variables were log-transformed. Linear regression analyses were then carried out (**Supplementary Table 4-6**) with a **Benjamini-Hochberg** correction for false-positive rate. Statistical significance was set at p < 0.05 after corrections for multiple comparisons. All non-significant results are noted NS and are all detailed in **Supplementary Table 4-7**.

Given that for the first two objectives of this paper (i.e. predicting adult behaviour based on adolescent PCA scores, and the links between adult behaviour and adult neurobiology) we analysed without assumptions, we applied a multiple comparison correction using the **Benjamini-Hochberg** p-value correction to limit false-positives (**Supplementary Table 4-6**). This allowed us to select the parameters to be used to assess the prediction between adolescent PCA scores and adult neurobiology by taking only variables that had already been correlated previously. Thus, for our final objective, we tested a hypothesis with a priori, without multiple comparisons, which allowed us to maintain a significance threshold of p < 0.05 (**Supplementary Table 7**).

#### Behavioural analysis

To see prediction from early behaviour with adult neurobiology, we looked sequentially at the links between adolescent PCA scores with adult behaviours, and then to the links between adult behaviours with adult metabolites and mRNA levels. To filter these data, we first selected only the variables significantly correlated between adolescent PCA scores and adult behaviours (**Figure 3**) to thereafter see which adult behaviours were correlated to adult neurobiological features (**Figure 4**). And finally, we have only selected the variables that were also correlated between adult behaviours and adult neurobiological features, to see which ones of these variables were correlated between adolescent PCA scores and adult neurobiological features (**Figure 5**).

#### Machine Learning analysis

After collection of all the raw behavioural data, they were first transformed into either a score (sucrose preference), a ratio (3-chambers task, looming task) or a percentage (social interaction task, 3-chambers task, looming task).

Then, the transformed data were normalised and centred to carry out Machine Learning analyses (PCA and HCPC, see below) using R.

Beforehand, variables that were too correlated with each other and co-dependent were excluded, to avoid giving them too much weight and having misleading redundancy in their construction.

To properly conduct these analyses, we chose a limited number of social variables to avoid having more behavioural variables compared to the sample size.

When conducting social tasks, a lot of different behaviours would be expressed and a large proportion of them would be sporadic and only expressed by some individuals only (i.e. communication, aggressive behaviour). However, to obtain social ‘profiles’, only variables expressed by all individuals were used, to ensure that they have a common baseline from which to compare their social abilities (i.e. SIT, 3-ch).

In addition, the effect of sex and environment was not addressed in this study, as it is fairly expected (and documented in the literature) to have biological differences based on these factors. In fact, the aim of our study isn’t to exclude this natural biological influence, but rather to explore whether we can go above these factors and explain the social profile by parameters other than sex and environment.

##### Principal Component Analysis (PCA)

The PCA is an unsupervised and well-known method used for dimensionality reduction of the data (in Forkosh et al., 2019). This analysis aim is to seek the variables association (into dimensions) that will maximise the variance of the data, and is extremely useful when working with data sets that have a lot of features such as behavioural data. Thus this analysis was used to summarise the social variables information in our data, by reducing the dimensionality of our sample without losing important information.

##### Hierarchical Clustering on Principal Components (HCPC)

Following the PCA analysis, a clustering method was used on the PCA output to identify specific groups according to their social dimensions. This method uses PCA coordinates of each individual and performs a dendogram to compute the distance between each individual. It allows us to define the optimal number of groups, the individuals who constituted them and the variables that differentiated them.

##### Linear Regression Models

In order to predict from adolescent social profile, both adult social behaviour and neurobiological features, some linear regression models were conducted in RStudio.

### Data Availability

The datasets generated during the current study are available from the corresponding author upon request.

### 2.5. Biological measures

The animals that were used to conduct the social behaviour analyses were also used to perform gene expression analysis by RT qPCR and basal brain metabolite levels by HPLC method (see below). Animals were killed by cervical dislocation right after isoflurane gas anaesthesia. Striatum, Olfactive Bulbs (OB), Cerebellum and PreFrontal Cortex (PFC) were dissected under binocular control and stored at −80 ◦C.

#### Microfluidic RT qPCR

##### RNA extraction

Total RNA was isolated and purified using a miRNeasy kit with Qiazol (QIAGEN reference 217004) according to the manufacturer’s instructions. Briefly, each tissue was shredded on ice for up to 2min at 20Hz using 5mm diameter steel beads (Stainless Steel Beads 5 mm x200, ref 69989) and a TIssuLyser II (Qiagen). Homogenates were then centrifuged at 4°C for 5 minutes at 12,000g. MaXtract High Density tubes (n. 129053, Qiagen) were then used to collect the aqueous phase.

Total RNA concentrations were assessed using Quant-IT RiboGreen (Invitrogen) and quality was controlled using a 4200 TapeStation system and RNA ScreenTape assays (Agilent Technologies Inc., Santa Clara, CA, U.S.A.). Only RNA samples with a RNA integrity number greater than 7 and a 18S/28S ratio greater than 1.7 were selected for further analyses.

##### Microfluidic RT qPCR

The RT was carried out with the High-Capacity cDNA Reverse Transcription Kits (n. 4368814, Thermofisher Scientific) without the DNase inhibitor, after contamination assessment. RT was conducted following the manufacturer’s instructions with minor modifications. As some samples had a low RNA concentration, to ensure a cDNA concentration of 500 ng/μL for all the samples, the following components volume were used to prepare the RT master mix: 2.35μL Nuclease free H²0, 3.5uL 10X RT buffer, 1.4μL 25X dNTP Mix (100 mM), 3.5μL 10X RT Random Primers and 1.75μL MultiScribe Reverse Transcriptase. This RT master mix provided a total solution of 12.5μL, which added to 22.5μL of RNA, leading to 35μL of cDNA. The final products were diluted in 20μL of Nuclease-free water and 55μL of TaqMan Master Mix (TaqMan Fast Advanced Master Mix) to a final concentration of 500 ng/μL of cDNA.

Microfluidic Real-time qPCR was performed with TaqMan Array Card using pre-filled TaqMan primers: dopamine transporter DAT (*Slc6a3*), dopamine receptor 2 *Drd2*, dopamine receptor 1 *Drd1*, serotonin (5-HT) receptor 1a *Htr1a*, serotonin (5-HT) receptor 2a *Htr2a*, serotonin (5-HT) receptor 2c *Htr2c*, serotonin (5-HT) transporter 5-HTT (*Slc6a4*), cholinergic nicotinic receptor beta 2 *Chrnb2*, cholinergic nicotinic receptor alpha 7 *Chrna7*, brain derived neurotrophic factor *Bdnf*, *18S*, hypoxanthine guanine phosphoribosyl transferase *Hprt*, nerve growth factor *Ngf*, cyclin-dependent kinase 5 *Cdk5*, cyclin D1 *Ccnd1*, oxytocin receptor *Oxtr*, arginine vasopressin receptor 1A *Avpr1a*, indoleamine 2,3-dioxygenase 1 *Ido1*, kynurenine 3-monooxygenase *Kmo*, tryptophan 2,3-dioxygenase *Tdo2*, kynureninase *Kynu*, glucuronidase beat *Gusb*, aryl-hydrocarbon receptor *Ahr*, nicotinamide phosphoribosyltransferase Nampt.

TaqMan Array cards (n. 4342249, ThermoFisher Scientific) were filled with 500 ng of cDNA following the manufacturer’s instructions. All the RT qPCR results are more detailed in **Supplementary Table 2**.

For each gene, common threshold fluorescence for all the samples was set into the exponential phase of the amplification and determined the cycle threshold (CT), corresponding to the number of amplification cycles needed to reach this threshold. All reactions were performed in duplicate and the mean value of CT was used for subsequent analysis. Relative gene expression quantification was performed using two endogenous housekeeping genes encoding the hypoxanthine phosphoribosyltransferase 1 (*Hprt1*) and the glucuronidase beta (*Gusb*).

##### Control gene

To select the control gene for each target gene by structure, we chose the control gene whose average CT was closest to the target gene (**Table 9** below). For most target genes, the closest control gene was *Gusb*. Another internal control gene, *18S*, was also used to control for quality and inter-plate variance.

**Table 9.**
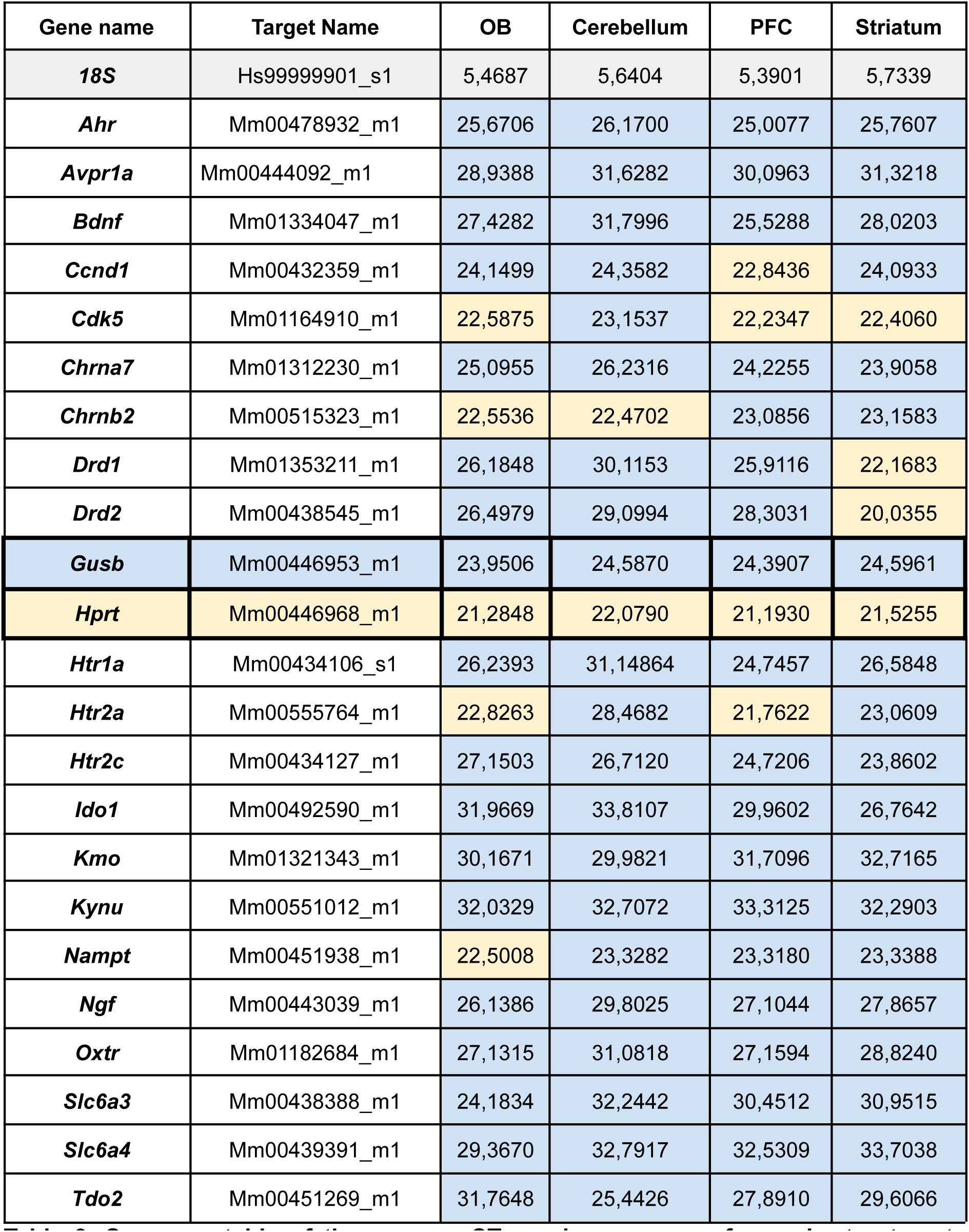
Summary table of the average CT number per gene for each structure to select the control gene. The control gene is selected based on the control gene whose average CT is closest to the target gene. For example, genes coloured blue have *Gusb* as their control gene while genes coloured yellow have *Hprt*.

**Table 10.** Optimized MS/MS parameters for the analysis of AccQ derivatized compounds, with the exception of underivatized adenosine.

| Compound | Precursor<br>(m/z) | DP<br>(V) | EP<br>(V) | Product Ion |  |  |  |  |  | IS |
| --- | --- | --- | --- | --- | --- | --- | --- | --- | --- | --- |
|  |  |  |  | Quantifier |  |  | Qualifier |  |  |  |
|  |  |  |  | Masse<br>(Da) | CE<br>(V) | CXP<br>(V) | Masse<br>(Da) | CE<br>(V) | CXP<br>(V) |  |
| Adenosine | 268.2 | 50 | 10 | 136.1 | 23 | 16 | 119 | 63 | 14 | 1 |
| Acide γ-amino-butyrique<br>(GABA) | 274.1 | 125 | 10 | 171 | 27 | 20 | 116 | 67 | 12 | 2 |
| Tyramine | 308.4 | 100 | 10 | 171.1 | 31 | 18 | 116.1 | 73 | 14 | 3 |
| Glutamate (Glu) | 318.1 | 95 | 10 | 171.1 | 29 | 20 | 116 | 75 | 14 | 4 |
| Glutamine (Gln) |  |  |  |  |  |  |  |  |  |  |
| Dopamine HCl (Dopa) | 324.2 | 150 | 10 | 171.1 | 31 | 20 | 116 | 77 | 12 | 5 |
| Norépinephrine (Norepi) | 340.2 | 145 | 10 | 171.1 | 31 | 20 | 116 | 79 | 12 | 6 |
| Serotonin | 347.2 | 120 | 10 | 171.1 | 35 | 20 | 160.1 | 27 | 18 | 7 |
| Tyrosine | 352.2 | 85 | 10 | 171.1 | 29 | 20 | 116 | 83 | 12 | 10 |
| Epinephrine (Epi) | 354.2 | 125 | 10 | 171.1 | 33 | 20 | 166.1 | 25 | 20 | 8 |
| Normetanephrine<br>(Normeta) | 354.2 | 125 | 10 | 171.1 | 33 | 20 | 166.1 | 25 | 20 | 9 |
| Tryptophane (Trp) | 375.1 | 155 | 10 | 171 | 31 | 20 | 116 | 87 | 12 | 10 |
| Kynurenine (Kyn) | 379.1 | 158 | 10 | 171.1 | 29 | 20 | 145.2 | 23 | 16 | 10 |
| 5-Hydroxytryptophane<br>(5-HTP) | 391.2 | 161 | 10 | 171.1 | 33 | 20 | 116.1 | 93 | 14 | 10 |
| IS 1 : Adenosine<br>13C10 15N5 | 283.2 | 50 | 10 | 146.1 | 23 | 16 | 129.1 | 63 | 14 |  |
| IS 2 : Acide<br>γ-amino-butyrique d6 | 280.3 | 125 | 10 | 171.0 | 27 | 20 | 116.0 | 67 | 12 |  |
| IS 3 : Tyramine d4 | 312.4 | 100 | 10 | 171.1 | 31 | 18 | 116.1 | 73 | 14 |  |
| IS 4 : Glutamate d5 | 323.2 | 95 | 10 | 171.1 | 29 | 20 | 116.0 | 77 | 12 |  |
| IS 5 : Dopamine d4 | 328.3 | 150 | 10 | 171.1 | 31 | 20 | 116.0 | 77 | 12 |  |
| IS 6 : Norepinephrine d6 | 346.2 | 96 | 10 | 171.1 | 33 | 20 | 116.0 | 81 | 12 |  |
| IS 7 : Serotonin d4 | 351.2 | 120 | 10 | 171.1 | 35 | 20 | 160.1 | 27 | 18 |  |
| IS 8 : Epinephrine d6 | 360.2 | 61 | 10 | 171.1 | 33 | 20 | 342.3 | 13 | 16 |  |
| IS 9 : Normetanephrine<br>d3 | 357.2 | 95 | 10 | 171.1 | 33 | 20 | 116.0 | 83 | 12 |  |
| IS 10 :<br>1-méthyl-Tryptophane | 389.2 | 155 | 10 | 171.0 | 31 | 20 | 116.0 | 87 | 12 |  |

#### HPLC: Basal Monoamine Brain Level Analysis

We studied in social brain areas (i.e. prefrontal cortex, olfactory bulb, striatum and cerebellum) different neurotransmission systems, such as the monoaminergic system (dopamine & serotonin), their precursors (Tryptophan, Tyrosine & 5-HTP) and their ‘degradation/synthesis’ product (noradrenaline, normetanephrine). This enabled us to make ratio of synthesis or degradation of the neurotransmitter for serotonin (Tryptophan/5-HTP, 5-HTP/5-HT), dopamine (Tyrosine/Dopamine) and noradrenaline (Dopamine/Noradrenaline & Noradrenaline/Normetanephrine). We also targeted trace-amines because they have an influence on social behaviour, and we especially studied the level of Tyramine as its synthesis ratio (Tyrosine/Tyramine). We also investigated the kynurenine pathway to look for the Kynurenine synthesis (Tyrosine/Kynurenine) or neuroprotective/neurotoxicity balance (Kynurenine/5HT). We also assessed the excitatory/inhibitory activity in the brain, by measuring levels of GABA & Glutamate, as the balance between them (GABA/Glutamate) and the degradation of Glutamate (Glutamate/Glutamine), but we also looked at Adenosine level that can also act as a neuromodulator of brain excitation/inhibition. All metabolites and turnover or synthesis levels are more detailed in **Supplementary Table 3.**

##### Chemicals

All chemical standards were purchased from Sigma Aldrich (Saint-Quentin-Fallavier, France), except for tyramine and serotonin d4 which were purchased from Santa Cruz Biotechnology (Heidelberg, Germany). LC–MS hypergrade acetonitrile (ACN), methanol (MeOH), chloroforme (CHCl3), LC–MS grade formic acid (FA) and 5-sulfosalicylic acid (SSA) were provided by VWR Chemicals (Rosny-sous-bois, France). Ultrapure water (UPW) was obtained from a Milli-Q® IQ 7003 water purification system (Merck Millipore, Guyancourt, France).

##### Sample preparation

Samples were exactly weighted in lysing CK14 tube and ground with a Precellys-Cryolys system (Bertin Technologies) as following: 3 cycles at 5,500 Hz during 10 seconds separated with a 10 seconds pause time. Grinding was carried out in MeOH containing an adapted concentration of each labelled internal standards (IS). A liquid-liquid extraction step based on Folch’s procedure was performed by adding volumes of cold extraction solvents adapted for each sample on a basis of CHCl3/MeOH/UPW ratios of 8/6/16 µL for 1 mg of tissue. After shaking for 5 minutes and then centrifugation (10 min, 18 000 g, 4°C), aqueous upper phase was transferred and dried under nitrogen flow at 30°C for 45 min. Residual was solubilized in 50 µL of UPW containing 0.05 % of FA. A precipitation step was then performed on 10 µL of sample with 10 µL of SSA (10% in UPW) and 10 µL of UPW and the supernatant was finally derivatized with AccQ-Fluor Reagent Kit (Waters, Guyancourt, France) before transferred in vial and injection in the LC-MS/MS system.

##### Liquid chromatography-tandem mass spectrometry

The liquid chromatography system consisted of an Exion LC coupled to a 6500+ QTRAP mass spectrometer from Sciex (Villebon-sur-Yvette, France) equipped with an electrospray source operating in positive ionization. The MS parameters were as follows: Curtain gas at 30 psi, Collision Activated Dissociation gas (CAD) at 6 psi (Medium), IonSpray Voltage at 4500 V, Source Temperature at 450°C, Ion source gas 1 (nebulizer gas) at 60 psi, Ion source gas 2 (desolvation gas) at 60 psi, Gas interface heater ON, resolution unit. MRM transitions, collision energies (CE) and fragmentor voltages for all compounds are given in table X. Chromatographic separation was performed on a Kinetex® Biphenyl column (2.1 mm i.d. × 150 mm, 1.7 μm, Phenomenex, le Pecq). Column and autosampler were maintained at 40 °C and 4 °C, respectively. The mobiles phases were composed of phase A (UPW containing 0.1% of FA) and phase B (MeOH containing 0.1% of FA). The flow rate was 0.4 mL/min. The gradient was as follow: 0 % B (3 min), linear increase in 4 min to 30 % B, linear increase in 9 min to 65 % B, linear increase in 1.5 min to 100 % B, held during 2 min and linear decrease in 0.5 min to 0 % B. A post-time re-equilibration of 8 min was applied. Injection volume was 2 µL and the injection system (needle, pump, port) was carefully washed to minimize carryover effect.

Acquisition software Analyst version 1.7 was used to control the equipment during the assay, and qualitative data processing was done with Sciex OS-Q software.

Analytes peak areas were normalized to those of the IS. As the extraction protocol has been standardized according to sample weight, peak area ratios were expressed per mg of tissue allowing a relative comparison of results between arms of subject.

## Acknowledgments

Plateforme IRBA, Véronique et Nicolas IRBA, Hélène Font This work was supported by the *Centre National de la Recherche Scientifique*, Paris-Saclay University, the *Agence Nationale de la Recherche* -ANR-Cerbot and was funded by the doctoral school BioSigne ED958. This research received support from IReSP and INCa as part of a call for applications for doctoral grants launched in 2024”: project “CADDOC24-363397”.

## Supplementary data

**Supplementary Figure 1.**
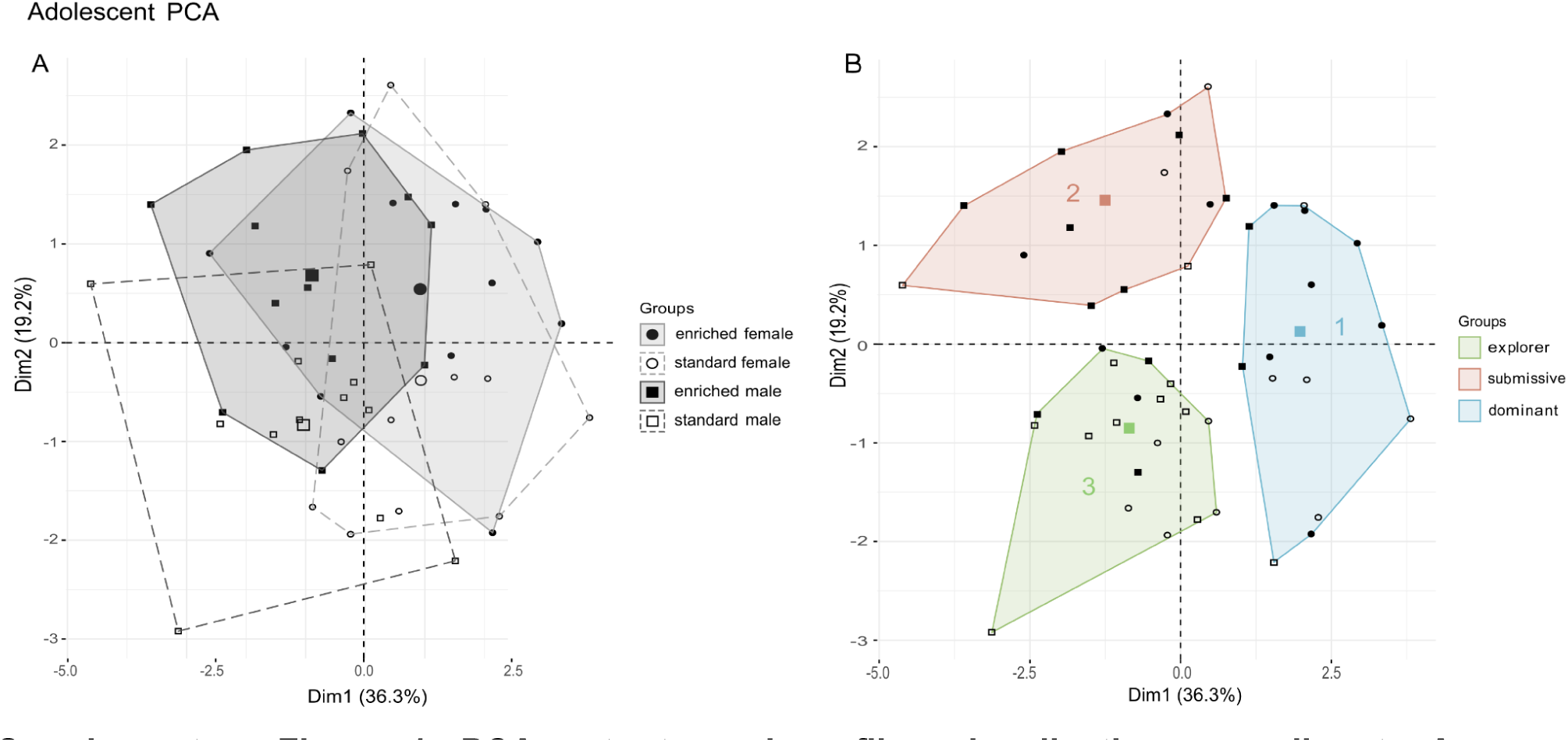
PCA outputs and profiles visualisation according to A. independent factors (sex, life-environment), or B. Hierarchical Clustering on Principal Components outputs for adolescent social behaviours. Results show that sex and environment factors **(A)** do not explain alone the variance of adolescent social behaviours as they display equal variance between them (Levene’s test: PC1: F(3, 44) = 0.2445, p = 0.8648; PC2: F(3, 44) = 0.3943, p = 0.7583; PC3: F(3, 44) = 0.8425, p = 0.4779). On the other hand, adolescent social profiles do explain better the variance in social behaviours compared to sex and environment factors, as their variance statistically differ between them (Levene’s test: PC1: F(2, 45) = 3.7966, p = 0.0299; PC2: F(2, 45) = 3.3313, p = 0.0447; PC3: F(2, 45) = 0.6878, p = 0.5078). All results are detailed in **Supplementary Table 1**.

**Supplementary Figure 2.**
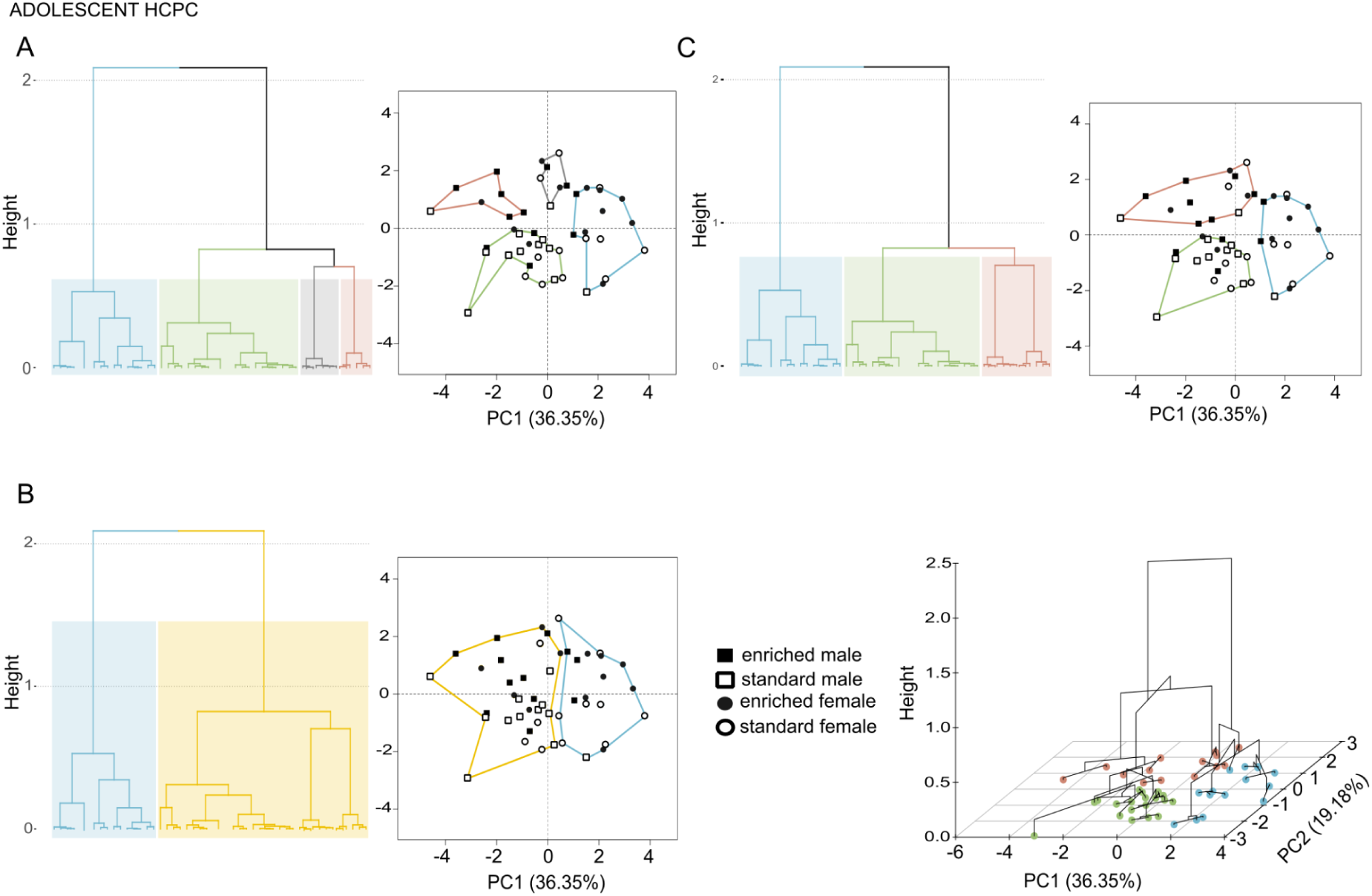
HCPC clustering analysis to compare and select the optimal number of social profiles on adolescent data. **A.** With 4 groups, **B.** 2 groups, and **C**. 3 groups. Depending on the choice threshold, we saw that it was more optimal to choose 3 clusters (C). Indeed, the variance seems to be better represented with 3 groups (C) as they statistically differ for PC1 (Levene’s test: PC1: F(2, 45) = 3.7966, p = 0.0299) and PC2 (Levene’s test: PC2 (F(2,45) = 3.3313, p = 0.0447), but not for PC3 unfortunately (F(2, 45) = 0.6878, p = 0.5079). While the other number of groups didn’t seem to be optimal to explain the variability as they didn’t differ statistically for each PCA score. For instance, 4 groups (A) wouldn’t represent the variability in social behaviours as they didn’t statistically differ for PC1 (Levene’s test: F(3, 44) = 0.7904, p = 0.5057), PC2 (Levene’s test: F(3, 44) = 2.0532, p = 0.1202) or PC3 (Levene’s test: F(3, 44) = 1.0351, p = 0.3863). It is also the case for choosing 2 groups, as they didn’t statistically differ for PC1 (Levene’s test: F(1, 46) = 0.8864, p = 0.3513), PC2 (Levene’s test: F(1, 46) = 0.0421, p = 0.8381) or PC3 (Levene’s test: F(1, 46) = 0.8100, p = 0.3727). All results are further detailed in **Supplementary Table 1**.

**Supplementary Figure 3.**
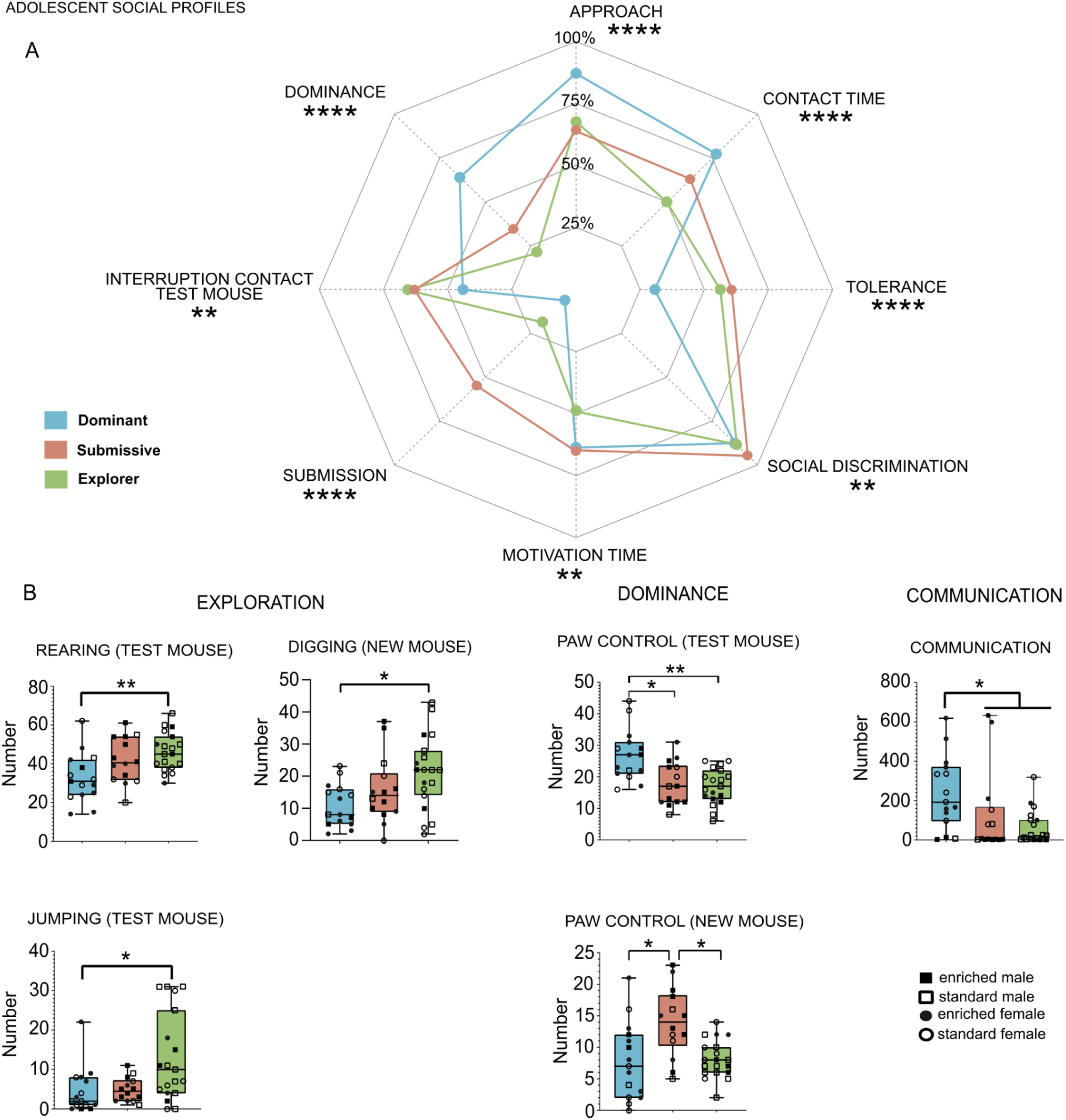
Statistical differences between social profiles for other social variables during interaction task in adolescence. **A.** Radarchart representing the contribution of social behaviours for each adolescent social profiles. **B.** Kruskal-Wallis analysis and Dunn’s pairwise comparison showed significant effects for: explorative behaviours (rearing (test mouse): dominant vs. explorer: p = 0.0025 ; digging (new mouse): dominant vs. explorer: p = 0.0075 ; jumping (test mouse): dominant vs. explorer: p = 0.0074), other dominant variables (paw control (test mouse): dominant vs. explorer: p = 0.0012 ; paw control (new mouse): dominant vs. submissive: p = 0.0051 ; explorer vs. submissive: p = 0.0078) and communication (dominant vs. explorer: p = 0.0091; dominant vs. submissive: p = 0.0152). * p < 0.05; ** p < 0.01; *** p < 0.001, **** p < 0.0001 ns not significant. All other social variables didn’t reach the significance threshold and are detailed in **Supplementary Table 1**.

**Supplementary Figure 4.**
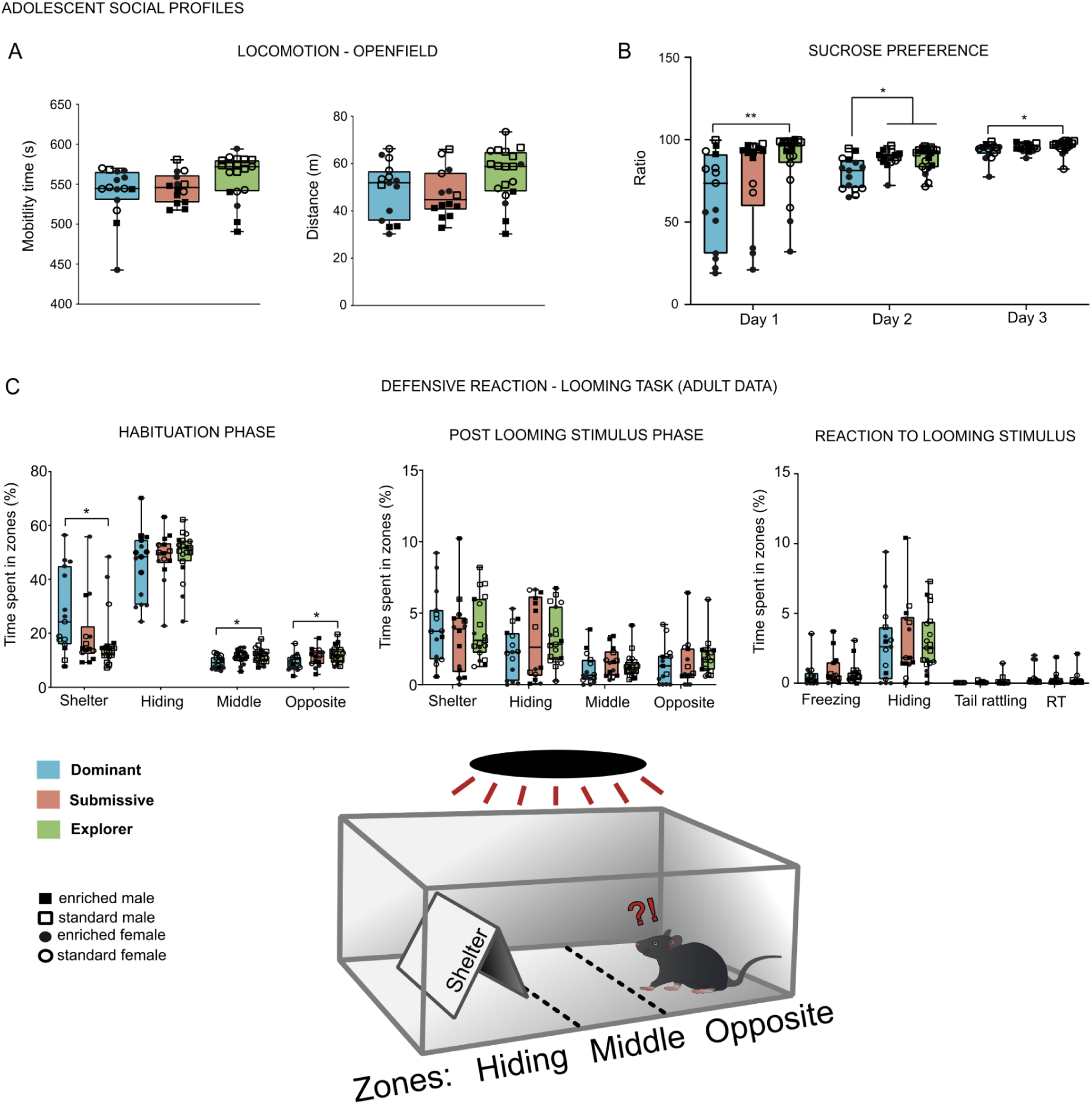
Statistical differences between social profiles for locomotion and sucrose preference in adolescence, and looming task in adulthood. **A.** Adolescent locomotion variables during Openfield between social profiles. The Kruskal-Wallis analysis showed no significant effects of locomotion (Mobility time: KW(2) = 5.981, ns; Distance: KW(2) = 4.740, ns) between the social profiles. **B.** Adolescent sucrose preference ratio between social profiles. The Kruskal-Wallis analysis showed significant differences of sucrose consumption during the first (KW(2) = 8.957, p = 0.0113), second (KW(2) = 9.220, p = 0.0099) and third days (KW(2) = 6.327, p = 0.0422) only between dominant individuals and explorer ones (Dunn’s test: Day1: p = 0.0046, Day2: p = 0.0087, Day3: p = 0.0240), but not with the submissive individuals, except on Day 2 (Dunn’s test: dominant vs. submissive: p = 0.0177). **C.** Adult looming task differences between adolescent social profiles. The Kruskal-Wallis analysis showed significant differences between profiles in time spent in the apparatus before the looming stimulus during habituation phase, especially under the shelter (KW(2) = 6.806, p = 0.0333), in the middle (KW(2) = 6.880, p = 0.0321) and in the opposite part of the apparatus (KW(2) = 7.965, p = 0.0186). Those differences were present between dominant individuals and explorers only (Dunn’s test: respectively p = 0.0156, p = 0.0212, p = 0.0076). Other variables of the looming task were not shown to be significantly different. * p < 0.05, ns not significant. All other non-significant results are detailed in **Supplementary Table 1**.

**Suppl. Fig. 5.**
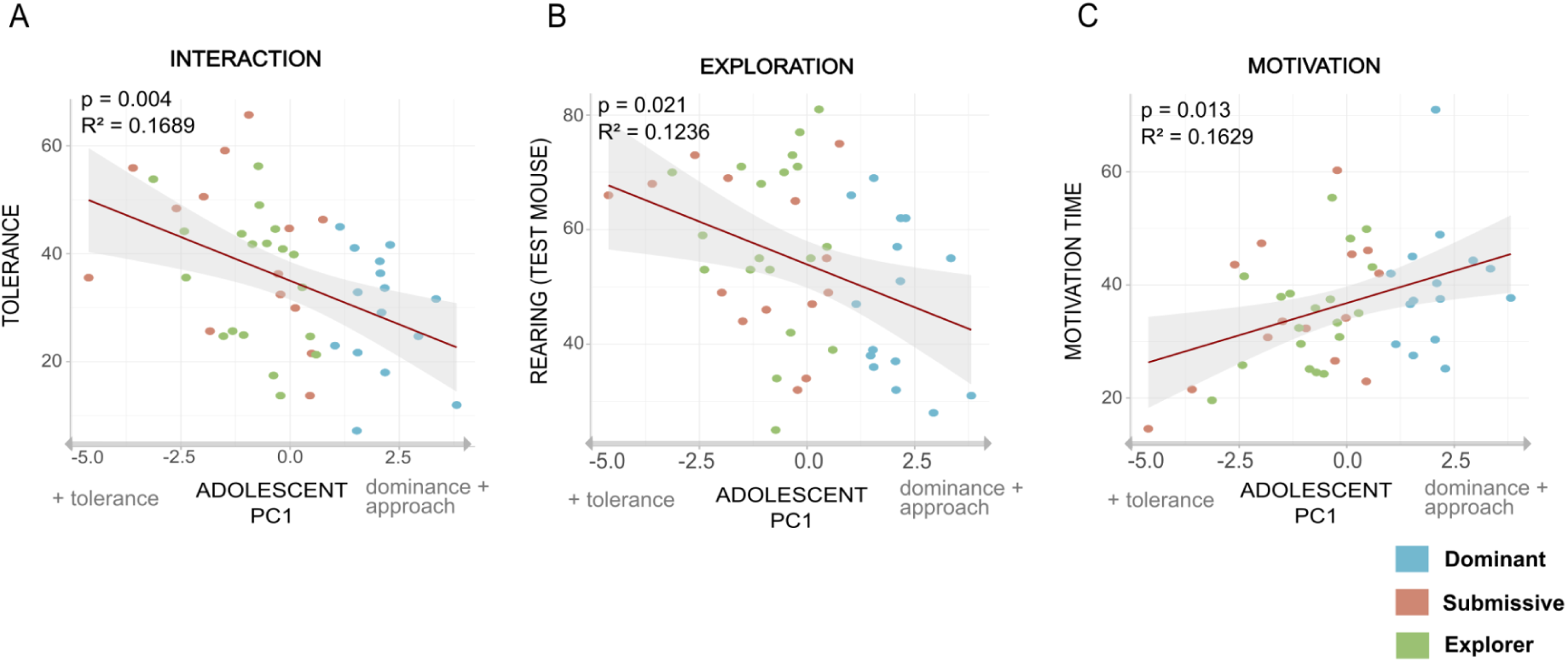
All prediction models from adolescent social behaviours of adult behaviours. Linear regressions between adolescent PCA scores and adult social behaviours for all social variables (used in PCA or not): **A.** dominance (dominance: F(3, 44) = 4.13, p = 0.002, R² = 0.1679; paw control (test mouse) x PC1: F(3, 44) = 6.716, p = 0.0048, R² = 0.2673; paw control (test mouse) x PC2: F(3, 44) = 6.716, p = 0.0091, R² = 0.2673) ; **B.** interaction (approach: F(3, 44) = 4.185, p = 0.004, R² = 0.1689; interruption of contact (test mouse): F(3, 44) = 2.89, p = 0.030, R² = 0.1077; tolerance: F(3, 44) = 4.185, p = 0.004, R² = 0.1689); **C.** exploration (jumping (test mouse): F(3, 44) = 4.326, p = 0.010, R² = 0.1751; rearing (test mouse): F(3, 44) = 3.209, p = 0.021, R² = 0.1236) ; **D.** grooming ((new mouse): F(3, 44) = 2.833, p = 0.0269, R² = 0.1048) and **E.** motivation (motivation time: F(3, 44) = 4.049, p = 0.013, R² = 0.1629). Each social profile is represented in colour, all non-significant results are detailed in **Supplementary Table 4**. Significant threshold : p < 0.05.

**Suppl. Fig. 6.**
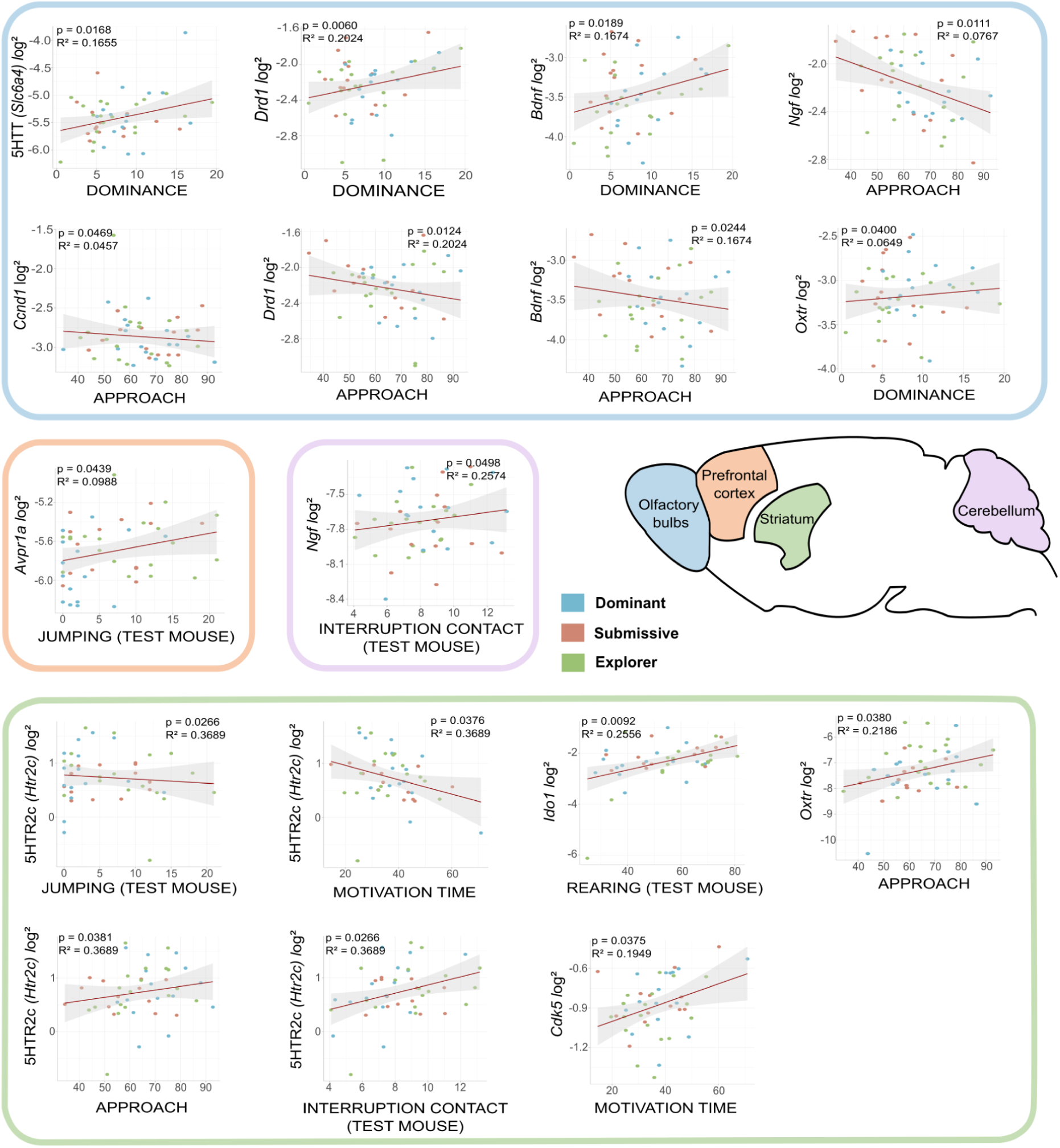
Adult social and non social behaviours predict adult basal brain gene expression. Linear regression models from adult social behaviours of RT qPCR levels in the social circuit. Results show significant predictions for 12 genes in the Olfactory bulbs (in blue), the Prefrontal cortex (in orange), the Striatum (in green) and the Cerebellum (in purple). All results are further detailed in **Supplementary Table 5** along with other results that didn’t reach the statistical corrected threshold.

**Suppl. Fig. 7.**
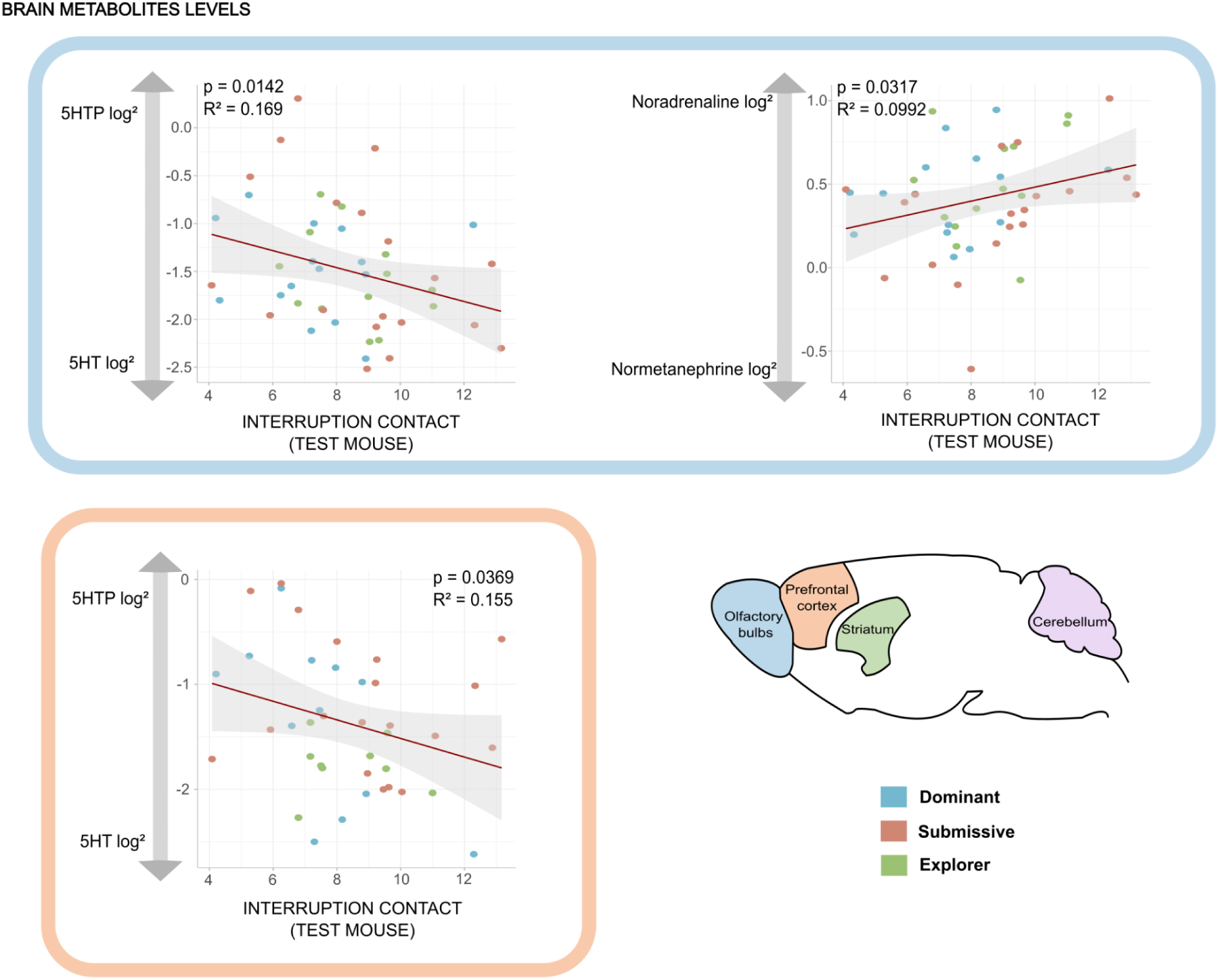
Adult social behaviours predict adult basal brain metabolites levels. Linear regression models from adult social behaviours of adult brain metabolites levels. Results shown are significant for **5HTP/5HT**, and **Noradrenaline/Normetanephrine** (**NA/Nor**) ratios in the Olfactory bulb (blue) and the Prefrontal cortex (orange). All results are detailed in **Supplementary Table 6** with other results that didn’t reach the statistical corrected threshold.

